# Hippocampal CA3 forms a two-layer network of molecularly distinct cell types in mice and humans

**DOI:** 10.64898/2026.07.04.736487

**Authors:** Rebecca J. Morse-Mora, Yoav Ben-Simon, Satish Arcot Jayaram, Karl Rössler, Peter Jonas, Jake F. Watson

## Abstract

Associational memory is critically dependent on the CA3 recurrent circuit, at the centre of the hippocampal system. While diversity amongst CA3 pyramidal neurons (PNs) has been long reported, the principles organising cell type identity and their consequences for circuit function remain unclear. Here, we demonstrate that deep-superficial is the primary axis of heterogeneity in CA3. PNs segregate into two molecularly distinct subtypes, superficial *St18−*, and deep *St18*+ neurons, with distinct anatomical, molecular, and functional properties. Using an *St18*-Cre mouse line, we reveal that these subtypes have divergent wiring patterns. Superficial neurons form the canonical broad autoassociative network across hemispheres and project to CA1, while deep *St18*+ PNs form a strikingly distinct, CA3-restricted, and predominantly unilateral projection — a previously unrecognised circuit module embedded within the hippocampal associative network. This anatomical and molecular organisation is conserved across mammals, including mice, pigs, and humans, where patch-clamp recording in human tissue demonstrates conserved functional hallmarks of each PN subtype. Together, these findings fundamentally revise the hippocampal CA3 circuit map, revealing it as two highly distinct circuit elements and providing the molecular tools to dissect the contribution of each layer to hippocampal computation and cognition.

## INTRODUCTION

Neuronal heterogeneity is a fundamental feature of brain organisation. Diverse morphological, physiological, and connective properties are thought to enable complex computational capabilities ^1^. Understanding neuronal diversity can unlock central principles of circuit computation ^2,3^, yet for many brain areas, the principal axes of heterogeneity are undefined. Without organisational clarity, diversity confounds rather than explains circuit function.

Hippocampal CA3 exemplifies this problem. The hippocampus is a central brain area for learning and memory ^4,5^, and its CA3 pyramidal neurons (PNs) form a broad, autoassociative network across hemispheres ^6–9^ — a circuit architecture that is uniquely suited for associational computation ^10–14^. CA3 PNs show diverse morphological ^15–19^, functional ^20–23^, and transcriptomic profiles ^24–26^, with differences following all three spatial axes (proximal-distal ^16,27–30^, dorso-ventral ^24,19,31,32^, and deep-superficial ^17,18,20,23,33^), as observed in other hippocampal areas ^34–40^. Transcriptomic analyses have identified between two and nine CA3 PN subtypes ^24–26^, yet without a molecular anchor unifying these observations, the organisational principles of CA3 PN heterogeneity have remained unresolved.

Here, we demonstrate that deep-superficial is the primary axis of heterogeneity in CA3, with PNs segregating into two molecularly distinct, fundamental subtypes. These subtypes are primarily distinguished by expression of the transcription factor *St18* (Suppression of Tumorigenicity 18), a genetic handle that allowed creation of a Cre-driver mouse line providing differential access to each subtype across the hippocampus. While superficial *St18−* PNs form the classical CA3 associational network, deep *St18+* cells have a highly distinct, non-canonical projection pattern. We show both molecular and functional conservation of deep PNs across mammals to humans, where we find conserved morpho-electric features in human hippocampal tissue. Together, these findings reframe how we consider the CA3 memory circuit, revealing it as not one heterogeneous population, but two highly distinct PN types with divergent contributions to the autoassociative network.

## RESULTS

### The deep-superficial axis defines CA3 PN identity

To approach CA3 neuronal heterogeneity in an unbiased and molecularly informed manner, we reanalysed publicly available single-cell RNA-seq datasets from the Allen Institute ^41^ (**Fig. 1**, **Supplementary Fig. 1**). After batch correction and dimensionality reduction, we performed graph-based clustering on all sequenced CA3 PNs. Even at the lowest tested resolution, CA3 PNs fell into two clear groups (Leiden resolution: 0.01; **Fig. 1a**). Cluster 1, comprising 71% of cells, was marked by expression of *Epha3*, *Nos1*, *Mdga1*, *Tafa1*, and *Ctxn2,* while cluster 2 expressed *St18*, *Gabrg3*, *Col6a1*, *Htr2c*, and *Cdh13* (**Supplementary Fig. 1c**). *St18* and *Col6a1* have been shown previously to have biased expression on the deep-superficial axis within the CA3 pyramidal cell layer ^24^. Mapping these clusters onto whole brain spatial transcriptomic datasets ^42,43^ showed that cluster 2 neurons occupied deep positions within the CA3 pyramidal cell layer across the dorso-ventral extent of the hippocampus, while cluster 1 neurons were consistently superficial (**Fig. 1b, Supplementary Fig. 1d**).

**Figure 1:**
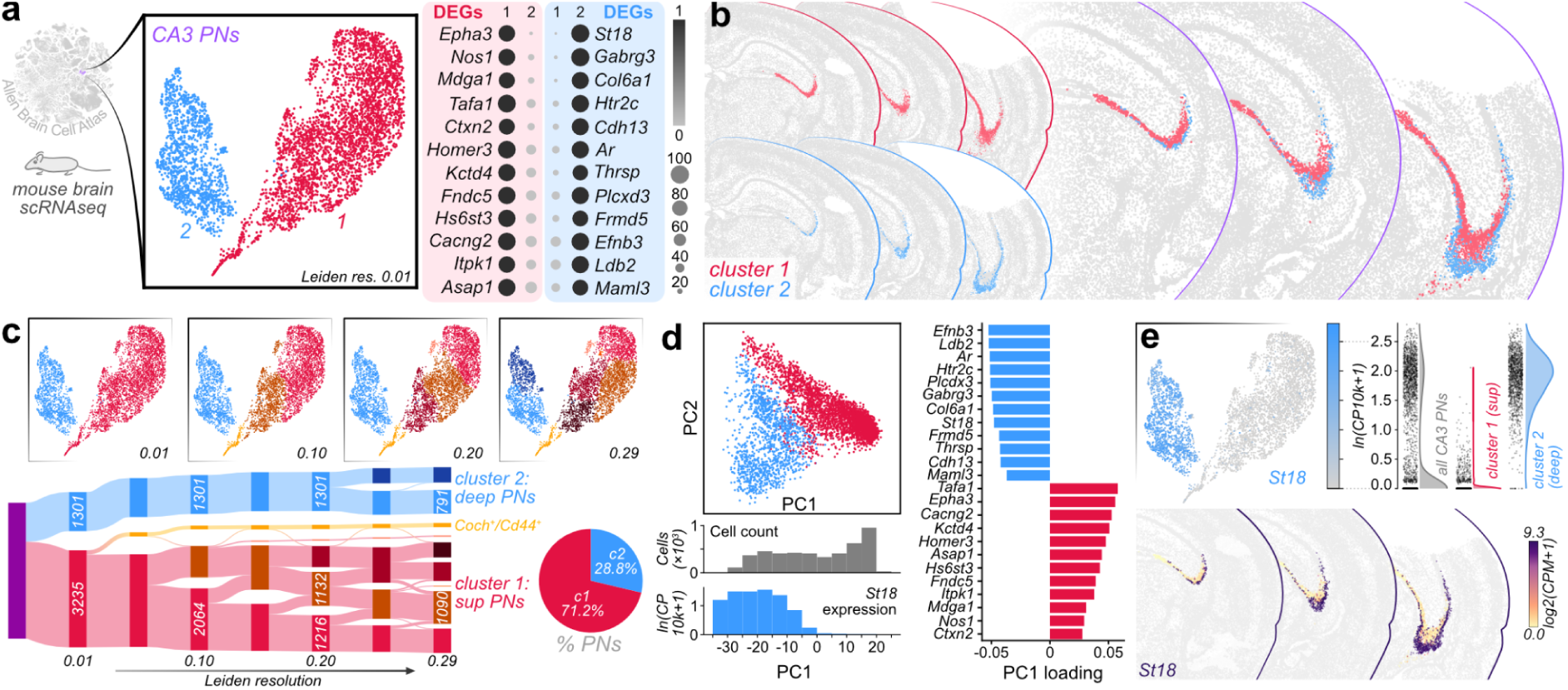
*St18*+ deep and *St18*− superficial PNs are the fundamental cell types of CA3. **a**, Transcriptomic analysis of CA3 PNs^43,41^ reveals two distinct genetic subtypes, presented in UMAP space (coloured by Leiden cluster). Dot plots of cluster-specific differentially expressed genes (DEGs) are coloured by the mean cluster expression (ln(CP10k + 1); where CP10k denotes counts per 10,000, per cell) scaled per gene (0–1), the dot size represents the fraction of cells expressing each gene within a cluster. **b**, Mapping clusters onto Allen Brain Cell Atlas spatial transcriptomic data shows that clusters 1 and 2 occupy superficial and deep positions within the CA3 pyramidal layer respectively, across the dorso-ventral and proximal-distal axes. Images presented individually (left) and merged (right; cropped to hippocampus). **c**, Increasing Leiden resolution shows interchanging subclusters within cluster 1 (superficial), suggestive of a heterogeneous population, depicted as a Sankey plot (nodes annotated with cell numbers). Deep (cluster 2) and superficial (cluster 1) cell types remain segregated across resolutions, consistent with stable cell types. A minor *Coch*+, *Cd44*+ subpopulation also shows stability from low resolutions (yellow). Pie chart shows proportions of CA3 PNs per subtype in this dataset. **d**, Principal component analysis (PC1 vs PC2) shows cluster identity following PC1. Histograms show cell density along PC1 (grey) and *St18* expression strongly polarised on this axis. All cluster-specific differentially expressed genes (from **a**) are similarly polarised along PC1. **e**, *St18* expression is highly restricted to cluster 2, confirmed by violin plots of expression per cluster. Spatial mapping onto the Allen Brain Cell Atlas validates *St18* expression in deep CA3 PNs (CPM denotes counts per million per cell).

The clarity of genetic cluster segregation suggested that deep and superficial neurons were distinct cell types. To test this hypothesis, we assessed the stability of graph-based cluster identity (**Fig. 1c**). At increasing resolution, the two primary clusters split into smaller groups. A minor group of *Coch*/*Cd44+* cells segregated within the superficial neuron cluster; a known subpopulation of CA3 PNs located in very ventral CA3 ^24,44^ (**Supplementary Fig. 1e**). Aside from this subgroup, the majority of clusters within the superficial neuron population were unstable, frequently splitting and merging as resolution increased (**Fig. 1c**). This pattern is consistent with a heterogeneous population organised along gradients rather than discrete boundaries. In stark contrast, no exchange occurred between primary clusters, confirming deep and superficial PNs to be distinct, stable cell types.

Next, we examined the principal components (PC) to better understand the variation driving cell identity (**Fig. 1d**). Cells followed a bimodal distribution along PC1, with cluster 2 (deep) and cluster 1 (superficial) neurons occupying distinct regions (**Fig. 1d**). Expression of *St18*, the most cluster-2-specific gene, was strongly polarised along PC1 (**Fig. 1d**), with other cluster-identifying genes (see **Fig. 1a**) loaded bidirectionally on PC1. This distribution confirms that deep-superficial identity maps to the first PC of CA3 PN heterogeneity (**Fig. 1d**). Previous studies have defined multiple subdivisions in CA3 PN types, laying out across both proximal-distal and dorso-ventral axes ^24^. Mapping these subdivisions onto our identified clusters showed each spatial region mapping within the superficial cluster. Heterogeneity was particularly evident on the proximal-distal axis, likely contributing to previously reported functional differences between CA3 PNs ^23,27,30^ (**Supplementary Fig. 1f**). These subdivisions however, did not map within the deep cluster, or across the deep-superficial distinction. Thus, deep and superficial PNs are distinct cell types, with superficial PNs forming a heterogeneous population with expression gradients following spatial locations.

Deep-superficial heterogeneity has been explored more extensively in CA1 ^34–36,45–47^. To compare the structure of PN heterogeneity between subfields, we applied equivalent analyses to CA1 PNs. Instead of two distinct subgroups, CA1 PNs formed one large population, with minor subpopulations segregating across clustering resolutions (**Supplementary Fig. 2**). Deep CA1 PNs are characterised by a *Col11a1+, Ndst4+*, *Calb1−, Syt17−* expression pattern ^48,49^. We could clearly identify the deep CA1 population on a transcriptomic map of CA1 PNs, yet it did not form a separate stable cluster as observed for deep CA3 PNs. Instead, deep CA1 PNs formed an unstable subcluster that appeared only at higher resolution, and interchanged cells with other clusters across resolution levels (**Supplementary Fig. 2**). These data are consistent with deep CA1 neurons forming part of a broader spectrum of heterogeneous CA1 PNs. Therefore, while the deep-superficial axis has genetically specialised PNs across CA regions, deep-superficial identity is a uniquely distinct cell type division in CA3.

Finally, we sought to identify a marker gene defining the deep CA3 PN cell type. *St18* was the gene with highest specificity for cluster 2 (**Fig. 1a**), and showed almost perfect segregation between clusters — highly expressed in virtually all deep PNs, and absent in superficial cells (**Fig. 1e**). Plotting *St18* expression on spatial transcriptomic data showed a correspondingly clear laminar distribution (**Fig. 1e**). Together therefore, CA3 comprises two fundamentally distinct PN types, deep *St18+* and superficial *St18−* neurons.

### Genetic access to deep PNs with *St18*-Cre mice

To gain genetic access to CA3 PN subtypes, we used CRISPR-Cas9 genome editing to introduce a T2A-iCre ^50^ element at the 3′ end of the *St18* coding region in C57BL/6 mice (**Fig. 2a**, referred to henceforth as *St18*-Cre mice). When crossed with an Ai9 reporter line (Cre-dependent tdTomato expression ^51^), we observed patterned tdTomato fluorescence throughout the mouse brain (**Fig. 2b**; **Supplementary Fig. 3**). In the hippocampus, we observed labelled cells across subfields, but clear labelling of PNs with deep somatic location in both dorsal and ventral CA3, consistent with the predicted identity of *St18+* deep PNs (**Fig. 2b**). We plotted the distribution of tdTomato+ PNs on both the proximal-distal (from the dentate gyrus (DG), towards CA2) and deep-superficial axes in coronal (dorsal) and horizontal (ventral) slices (**Fig. 2c**). While we observed labelled cells occupying more superficial positions, the vast majority of PNs had a somatic location at the border between *stratum pyramidale* and *stratum oriens* (**Fig. 2c,d**). This delineation was less apparent in proximal CA3, where labelled neurons were less abundant and more distributed in the radial axis (**Fig. 2c**). Both the varied distribution of *St18+* PNs and the transcriptomic differences within superficial cells support a model of proximal and distal CA3 as distinct subareas ^22,23^. We observed no differences suggestive of a further subdivision within CA3, therefore, ‘proximal’ and ‘distal’ terminology better captures CA3 architecture than CA3a, b, and c ^46^. In total, 21% of PNs in ventral CA3, and 12% in dorsal CA3 were tdTomato+, aligning closely with previous estimates for the abundance of deep PN populations ^17,18,23^ (**Fig. 2d**). Crucially, *St18+* and *St18−* neurons were labelled across the longitudinal axis of the hippocampus, confirming the two-layered architecture of CA3 as a fundamental organising principle for this brain area.

**Figure 2:**
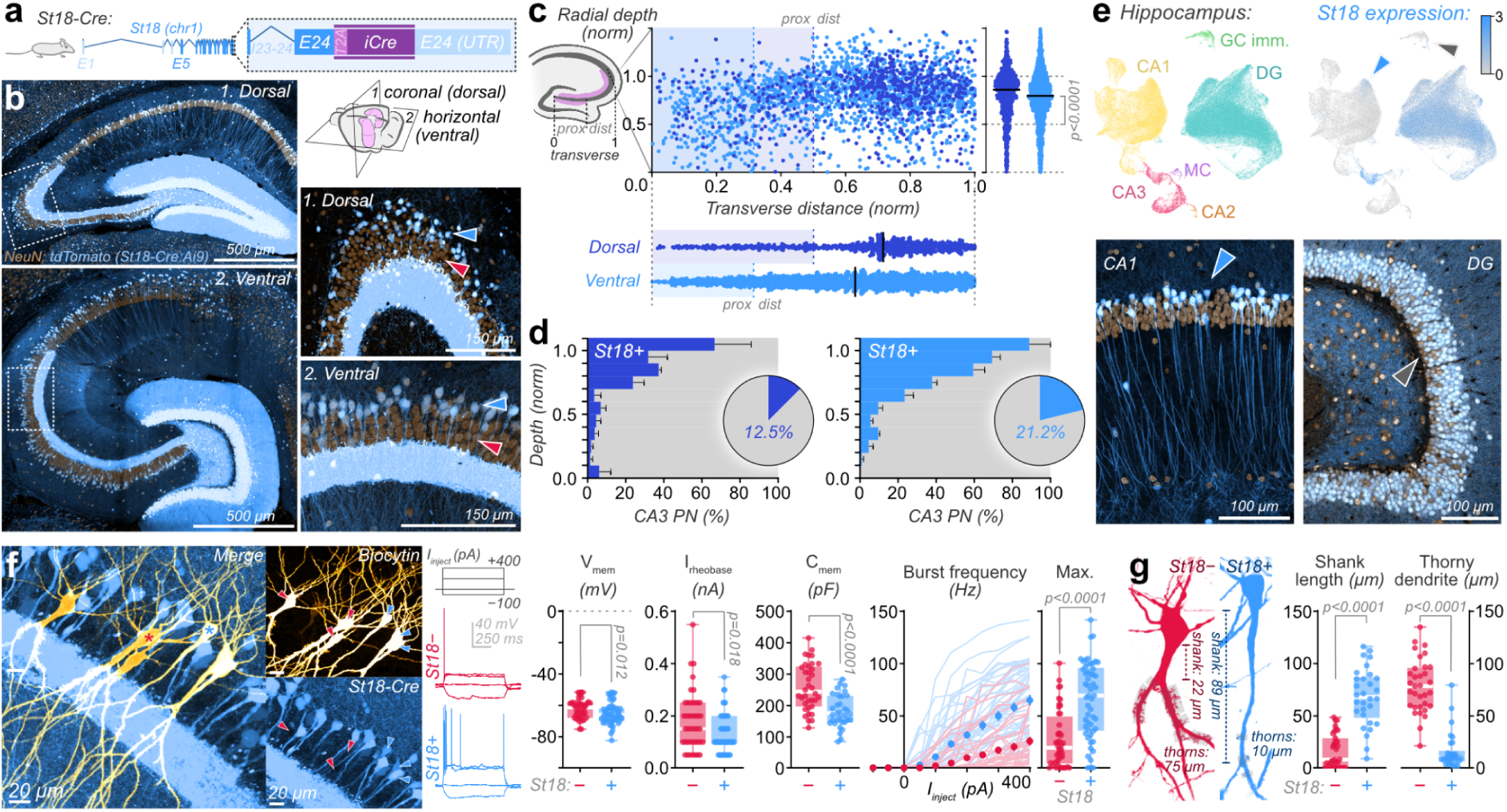
*St18*-Cre mice provide genetic access to deep CA3 PNs. **a**, *St18* genomic modification schematic: T2A-iCre was inserted into exon 24 (E24) after the protein coding region. **b**, Visualising *St18*-Cre expression with a tdTomato reporter crossing (Ai9) reveals specific laminar labelling of radially deep PNs across the proximal-distal and dorso-ventral axes (blue, tdTomato; brown, anti-NeuN; boxed area magnified; arrowheads indicate subtype layering). **c**, Quantification of *St18+* PN distribution confirmed their biased somatic location deep in the pyramidal cell layer, with preferential location in distal CA3 (median (interquartile range): *dorsal,* depth (norm) 0.86 (0.67, 0.98), transverse distance (norm) 0.72 (0.50, 0.83), *n* = 717; *ventral*, depth (norm) 0.80 (0.64, 0.95), transverse distance (norm) 0.63 (0.41, 0.81), *n* = 1684, *N* = 8 animals; one-sample Wilcoxon signed-rank test vs theoretical value 0.5, *p* < 0.0001; line represents median). **d**, 12.5% of PNs were *St18+* in dorsal, and 21.2% in ventral CA3, predominating in radially deep somatic locations (histogram: fraction of *St18+* neurons, mean ± SEM; *N* = 3 animals; *dorsal, St18− n* = 631, *St18*+ *n* = 90; *ventral, St18− n* = 636, *St18*+ *n* = 171). **e**, A subset of deep CA1 PNs (blue arrowhead), and mature DG GCs were also *St18+* in scRNA-seq data (Allen Brain Cell Atlas), correlating with tdTomato expression patterns in *St18*-Cre mice (GC: granule cells; MC: mossy cells; *St18* expression level in ln(CP10k+1)). **f**, Patch-clamp recorded neurons (biocytin filled; gold) from *St18*-Cre mice (blue) with example traces upon current injection. Arrows indicate *St18+* (blue) and *St18−* (red) cells, and traces are from cells marked *. Resting membrane potential (median (IQR): *St18−*, −61.2 mV (−67.4, −58.6), *n* = 44; *St18+*, −66.1 mV (−69.7, −62.1), *n* = 54; Mann-Whitney, *p* = 0.012), rheobase current (*St18−*, 150 pA (100, 250), *n* = 44; *St18+*, 100 pA (100, 200), *n* = 54; *p* = 0.018), and cell capacitance (*St18−*, 243 pF (199, 324), *n* = 36; *St18+*, 175 pF (147, 230), *n* = 34; *p* < 0.0001) were all lower in *St18+* than *St18−* neurons. Burst firing was more prominent in *St18+* neurons, evident both upon increasing current injection (symbols present mean ± SEM) and in maximal frequencies (Max.: *St18−*, 19.3 Hz (4.9, 49.4), *n* = 44; *St18+*, 69.4 Hz (36.4, 95.8), *n* = 54; *p* < 0.0001). **g**, *St18+* neurons show ‘long shank’ and ‘sparsely thorny’ morphological features (shank length: *St18−*, 8.0 μm (0.0, 28.7), *n* = 35; *St18+*, 68.2 μm (48.8, 87.0), *n* = 30; *p* < 0.0001. Thorny dendrite length: *St18−*, 78.8 μm (59.0, 96.2), *n* = 33; *St18+*, 9.8 μm (6.8, 16.6), *n* = 30; *p* < 0.0001).

In addition to CA3, we also observed labelling of neurons with deep somatic location in other CA subfields, and in the majority of DG granule cells (GCs) (**Fig. 2b**). We returned to the Allen Institute transcriptomic data to understand the observed expression pattern. Plotting *St18* expression across all hippocampal glutamatergic neurons recapitulated the clear segregation between CA3 PN types, while also demonstrating expression in a subset of CA1 PNs, and the majority of DG GCs (**Fig. 2e**). In *St18*-Cre mice, labelled CA1 neurons were located deep in the pyramidal cell layer, and *St18* RNA expression coincided with markers of deep CA1 PNs (**Fig. 2e**; **Supplementary Fig. 2**). Similarly, whilst the majority of DG GCs showed both *St18* expression in RNA-seq data and tdTomato labelling, immature GCs were unlabelled in RNA-seq data, which correlated with a lack of *St18* expression in the innermost region of the DG GC layer (**Fig. 2e**). In addition to hippocampal projection neurons, we also observed fluorescence in a subset of putative interneurons in *stratum oriens*, *radiatum*, and *lacunosum moleculare* (**Fig. 2b; Supplementary Fig. 3**), while outside the hippocampus, sparse neocortical neurons (layer 2-3 and 6b), olfactory bulb, and cerebellar granule neurons were labelled (**Supplementary Fig. 3**). In most of these cases, transcript expression patterns directly matched observed fluorescence (**Supplementary Fig. 3**). Some populations (e.g. L2-3, L6b) do not show *St18* RNA expression in adult mice ^43^, and cell labelling may therefore reflect transient *St18* expression during development, sufficient for long-term labelling with the Ai9 reporter line. While not exclusive to deep CA3 PNs, *St18*-Cre mice faithfully report expected *St18* expression patterns in RNA-seq datasets, and provide genetic access to distinct, transcriptomically identified principal neuron populations across the hippocampus.

We next sought to functionally and morphologically characterise tdTomato+ neurons in CA3. Previous work has shown that CA3 PNs with deeper somatic location have a bursting action potential (AP) phenotype ^52,33,21,17,23^, long primary dendritic shaft ^15,20^, and only sparse thorny excrescences ^15,17,21–23^. In patch-clamp recordings, *St18+* neurons showed significantly stronger burst firing than *St18−* cells, as well as a lower current threshold for AP generation, and lower cell capacitance (**Fig. 2f; Supplementary Fig. 3**). Morphologically, we observed characteristically long primary dendrites (**Fig. 2g**) with lower abundance of thorny excrescences on *St18+* cells (**Fig. 2g**), conferring greatly reduced input from the DG as has been previously reported ^15,17,22,18,23^. Therefore, the deep *St18*+ CA3 PN type unifies previously described ‘long shank’ ^15,22^, ‘athorny’ ^17^, ‘sparsely thorny’ ^20,22,18^, and ‘deep bursting’ neurons ^21,23^ as a defined molecular cell type.

### Developmental emergence of CA3 PN types

Developmental timing is a key determinant of neuronal heterogeneity. Neuronal layers are built sequentially from deep to superficial in both the neocortex ^53^ and CA1 ^54^, and neurons born at the same developmental timepoint are proposed to contribute similarly to circuit function and cognitive processing ^55,56^. In CA3, early-born neurons have deeper somatic locations, with distinct morphological and firing properties ^33^, suggesting that deep *St18+* and superficial *St18−* CA3 PNs may differentiate sequentially during development.

To test this hypothesis, we studied PNs born at different embryonic (E) timepoints (early-(E12.5), mid- (E14.5), and late-born (E16.5)), using *Ngn2*-CreER mice for birth-timed neuronal labelling (**Fig. 3**, **Supplementary Fig. 4**) ^33,57^. Neocortical and CA1 neurons showed sequential deep-to-superficial generation as expected, which was also observed in CA3, albeit with broader distributions (**Fig. 3a,b, Supplementary Fig. 4**). Patch-clamp recordings of birthdated CA3 PNs revealed no significant differences in burst firing or rheobase current, the hallmark electrophysiological features distinguishing deep and superficial CA3 PNs (**Fig. 3c,d**). Given the clear functional differences we observe between genetically identified PNs (**Fig. 2f**), these data suggest that developmental birthdate alone cannot predict the molecular cell type identity of CA3 PNs. To directly link developmental timing to molecular cell type identity, we performed bromodeoxyuridine (BrdU) birthdating in *St18*-Cre mice. BrdU labelling recapitulated the sequential laminar generation seen with *Ngn2*-CreER in CA3 and CA1 (**Fig. 3e**). *St18+* neurons were labelled by BrdU injections at all developmental timepoints, but with a decreasing likelihood from E12.5 to E16.5. However, only 17% of CA3 PNs labelled at E12.5 were *St18+*, confirming that molecular cell types are generated in an overlapping manner (**Fig. 3e**). Thus, generation of the two CA3 PN subtypes is temporally shifted during embryonic development, but not strictly defined. Genetic identity therefore provides the necessary lens to dissect their circuit contributions.

**Figure 3:**
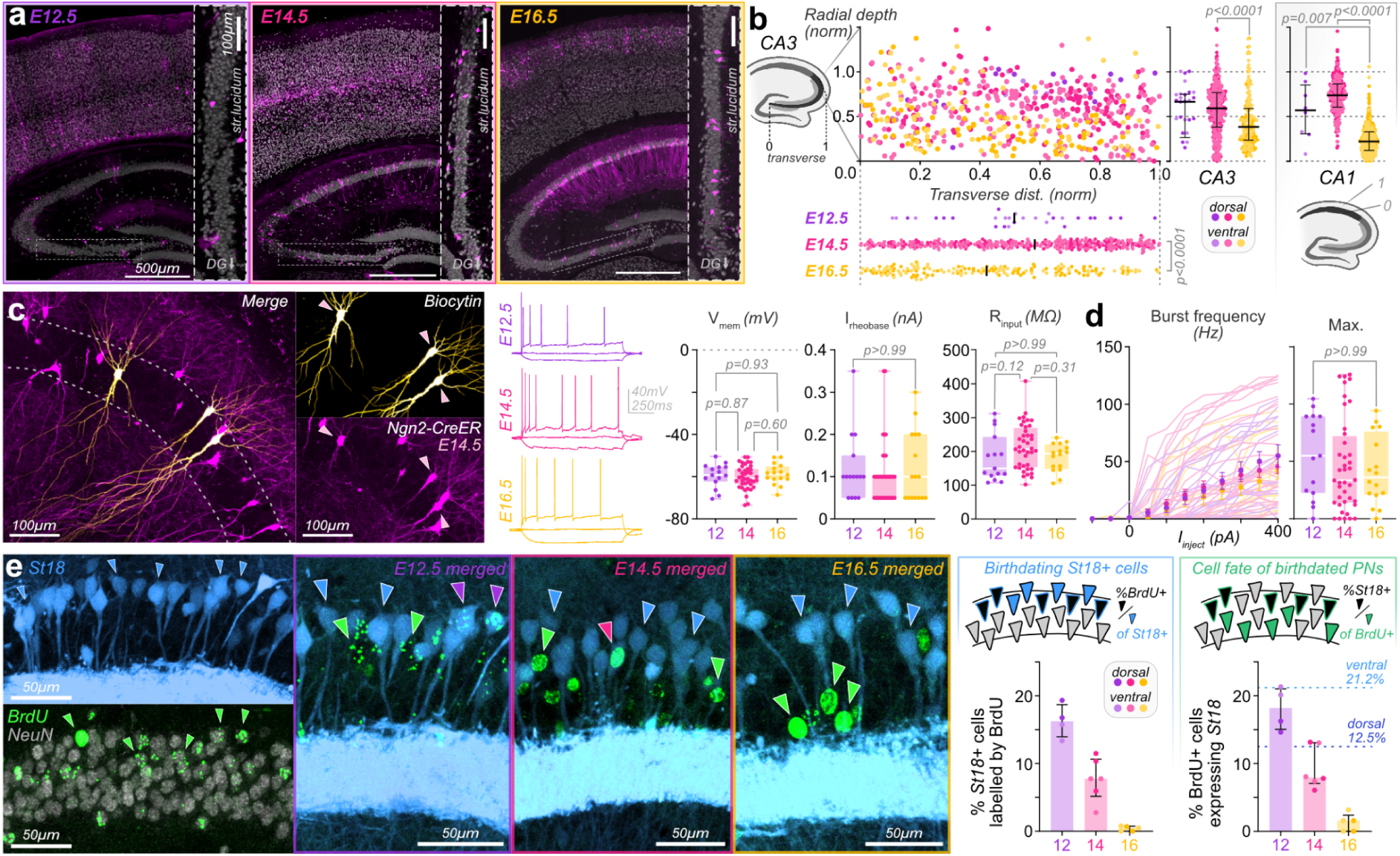
*St18*+ CA3 PNs are produced preferentially early in development. **a**, *Ngn2*-CreER:tdTomato labelling induced at embryonic timepoints, E12.5, E14.5, and E16.5 (magenta, tdTomato; grey, anti-NeuN; boxed areas magnified). **b**, Labelling shows more superficial and more proximal somatic location of later-born CA3 PNs (E16.5), with similar laminar distribution observed in CA1 (right) (median (IQR): depth (normalised); E12.5, 0.67 (0.26, 0.75), *n* = 29; E14.5, 0.59 (0.38, 0.77), *n* = 370; E16.5, 0.38 (0.23, 0.59), *n* = 16; Kruskal-Wallis, *p* < 0.0001. Transverse distance (normalised); E12.5, 0.51 (0.38, 0.68), *n* = 29; E14.5, 0.58 (0.31, 0.78), *n* = 370; E16.5, 0.42 (0.18, 0.67), *n* = 166; Kruskal-Wallis, *p* < 0.0001. CA1 depth (normalised): E12.5, 0.57 (0.31, 0.85), *n* = 8; E14.5, 0.74 (0.61, 0.86), *n* = 263; E16.5, 0.22 (0.12, 0.33), *n* = 716; Kruskal-Wallis, *p* < 0.0001; line represents median, and error bars denote IQR). CA1 layering was notably stricter than CA3, with a narrower distribution of labelled cells (CA3 vs CA1 IQR: E14.5, 0.39 vs. 0.26; E16.5, 0.35 vs. 0.21) (E12.5: *N* = 4 animals from 4 timed matings (TMs); E14.5: *N* = 3 animals, 3 TMs; E16.5: *N* = 3 animals, 3 TMs). **c**, Patch-clamp recording of embryonically labelled CA3 PNs (biocytin, gold; *Ngn2*-CreER tdTomato, magenta) showed no significant difference in functional properties, contrasting with differences observed in *St18*-labelled animals (median (IQR) V_m_: E12.5, −59.2 mV (−62.3, −55.4), *n* = 15 cells, *N* = 13 animals, 9 TMs; E14.5: −60.7 mV (−63.6, −56.3), *n* = 40, *N* = 14 animals, 8 TMs; E16.5: −58.7 mV (−61.1, −55.2), *n* = 16, *N* = 9 animals, 6 TMs; one-way ANOVA, *p* = 0.613. Rheobase current: E12.5: 100 pA (50, 150), *n* =15; E14.5, 100 pA (50, 100), *n* = 40; E16.5, 100 pA (50, 200), *n* = 16; Kruskal-Wallis, *p* = 0.647. Input resistance: E12.5, 149.7 MΩ (128.4, 242.8), *n* = 15; E14.5, 207.5 MΩ (154.2, 269.4), *n* = 40; E16.5, 193.8 MΩ (148.6, 220.5), *n* = 16; Kruskal-Wallis, *p* = 0.067). **d**, Burst frequencies upon current injection and maximum burst frequencies showed no embryonic timepoint-dependent difference. (Max. burst frequency: E12.5, 55 Hz (22.6, 89.3), *n* = 15; E14.5, 33.6 Hz (14.2, 72.1), *n* = 40; E16.5, 36 Hz (21, 75.4), *n* = 16; Kruskal-Wallis, *p* = 0.555). **e**, BrdU labelling at embryonic timepoints in *St18*-Cre mice (arrowheads indicate *St18*+ (blue), BrdU+ (green), and co-localized *St18*+, BrdU+ (pink) CA3 PNs). The percentage of *St18*+ CA3 PNs decreases across timepoints, indicating that these cells are earlier-born (%*St18*+/BrdU+; median (IQR) E12.5: 16.3% (14, 18.7), *n* = 4 slices, *N* = 2 animals, 2 TMs; E14.5: 7.8% (5.2, 10.6), *n* = 6 slices, *N* = 3 animals, 3 TMs; E16.5: 0.5% (0, 0.76), *n* = 5 slices, *N* = 3 animals, 3 TMs), but less than 20% of CA3 PNs born at E12.5 are *St18*+ (%BrdU+/*St18*+; E12.5: 18.2% (15, 21); E14.5: 8% (7, 13); E16.5: 1.6% (0, 2.4)).

### Non-canonical projections of deep CA3 PNs

Axonal projection patterns are a primary determinant of neuronal function, and were the basis for our current theories of hippocampal function ^10,12^. We therefore exploited the genetic access provided by *St18*-Cre mice to map the projection architecture of each CA3 PN subtype. We performed unilateral stereotactic co-injections of AAVs expressing orthogonal fluorescent reporters using either a ‘Cre-on’ Double-floxed, Inverted ORF (DIO) or a ‘Cre-off’ Single-floxed, Excisable ORF (SEO), simultaneously labelling deep and superficial CA3 PNs with tdTomato and EGFP, respectively (**Fig. 4a**). At the injection site in dorsal CA3, we observed the expected laminar labelling of deep and superficial CA3 PNs, with their differential abundance of thorny excrescences clearly evident (**Fig. 4a**). EGFP-labelled axons of superficial PNs showed the textbook projection pattern for CA3 neurons: strongly innervating both *stratum oriens* and *stratum radiatum* of both CA3 and CA1 in both ipsilateral and contralateral hemispheres. Their axons spread widely on the dorso-ventral axis, forming the broad recurrent network that is a central feature of CA3, and the canonical connection of the hippocampal ‘trisynaptic’ pathway to CA1 ^7,58^ (**Fig. 4b,c**). In contrast, deep PNs showed a strikingly different pattern. Three key features break with the classical projection pattern of CA3 PNs. First, we observed almost total confinement to the ipsilateral hemisphere. A weak projection from dorsal CA3 to *stratum oriens* of contralateral CA3 could be observed, but axons in the contralateral hemisphere were almost entirely absent from both CA1 and the mid-ventral hippocampus (**Fig. 4b–e**). Injections targeted both left and right hemispheres independently, each showing equivalent ipsilateral projections. This confirms that while deep PN projections are largely confined to each hemisphere, the overall circuit architecture is bilaterally symmetric. Previous single-neuron projection datasets classified CA3 PNs into ‘projection-types’, with around 75% of CA3 PNs projecting bilaterally, and 25% showing a unilateral intrahippocampal projection ^19^. Our data assign molecular identity to these projection-types, with both the proportions and projections approximately matching deep and superficial CA3 PNs. Second, deep PN projections had greatly reduced innervation of CA1. In both dorsal and ventral hippocampus, we observed far greater innervation of CA3 than CA1, with around five-fold lower CA1 innervation by deep-than by superficial PNs (**Fig. 4d**). Deep *St18+* PNs therefore contribute little to the primary output projection of CA3, instead acting predominantly *within* the CA3 associational network. Third, while longitudinal projections of superficial CA3 target the full extent of both *stratum oriens* and *radiatum*, *St18+* axons showed a specific innervation pattern at the very base of *stratum radiatum*, bordering *stratum lacunosum moleculare* (**Fig. 4b–e**). The site of this projection is abundant in interneuron somata ^59^, and recent data shows a specific LFP signal originating at this laminar border ^60^. Together therefore, while superficial *St18−* CA3 PNs form the canonical, textbook CA3 associative network and downstream projection to CA1 first described by Ramón y Cajal ^58,6^, deep *St18+* PNs have a highly distinct architecture, primarily innervating CA3 within but not between hemispheres, somewhat akin to an excitatory interneuron.

**Figure 4:**
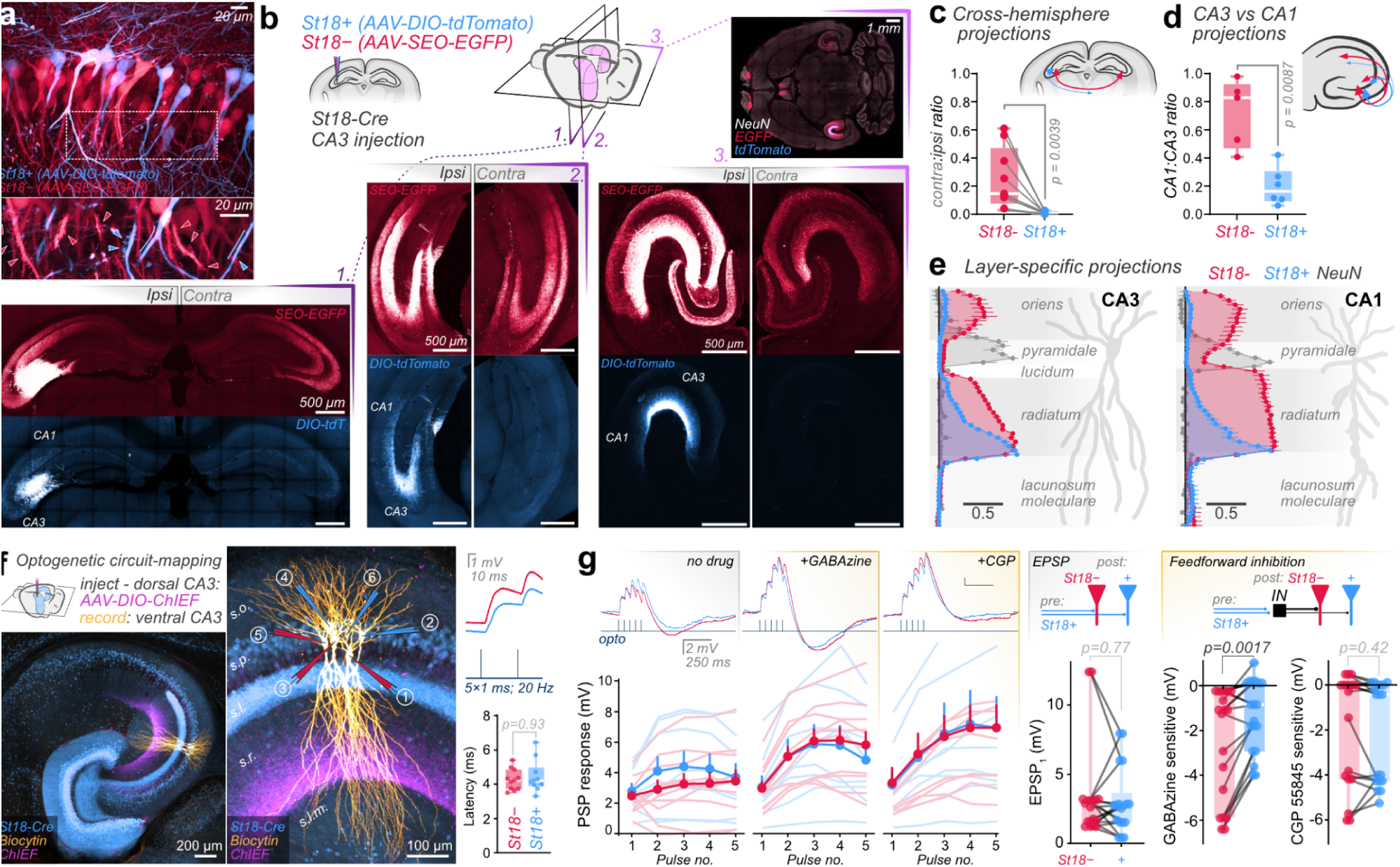
*St18*+ CA3 PNs show a non-canonical unilaterally targeted projection pattern. **a**, AAV-based labelling of *St18+*/*−* PNs. Inset: differential thorny excrescence abundance (arrowheads, thorns; line, lack thereof). **b**, Unilateral viral labelling shows hippocampal long-range projections of each subtype, depicted in three sections (1: coronal, bregma −1.9 mm; 2: bregma −2.7 mm; 3: horizontal). *St18−* PNs form the canonical CA3 projection, while *St18+* PNs form a specific and largely unilateral projection. **c**, Axon projections in the contra vs ipsilateral hemisphere were substantial for *St18−*, but minimal for *St18+* PNs (median (IQR); *St18−*: 0.15 (0.08, 0.47), *St18+* 0.01 (0.00, 0.02), *n* = 9 slices, *N* = 9 animals (6 coronal, 3 horizontal), Wilcoxon test, *p* = 0.0039). **d**, The CA1 vs CA3 projection ratio was far greater for *St18−* than *St18+* PNs (horizontal sections; *St18−*: 0.83 (0.47, 0.93), *N* = 5 animals; *St18+* 0.16 (0.10, 0.31), *N* = 6 animals; Mann-Whitney test, *p* = 0.0087). **e**, *St18−* PN projections were present in *stratum oriens* and *radiatum* of CA3 (left) and CA1 (right), while *St18+* axons projected specifically to the base of *stratum radiatum* (mean ± SEM, fluorescence normalised to peak, scale bar: 0.5 normalised units; CA3, *n* = 3 horizontal slice profiles; CA1, *n* = 9: 6 coronal, 3 horizontal). **f**, Optogenetic circuit-mapping from deep *St18+* in dorsal to ventral CA3 produced fast latency PSPs on a pair of *St18±* neurons (median (IQR) latency from stimulation onset: sup, 4.2 ms (3.8, 4.9), *n* = 11; deep, 4.2 ms (3.9, 5.0), *n* = 10; Mann-Whitney test, *p* = 0.93). **g**, Sequential application of GABAzine and CGP 55845 allowed dissection of excitatory and feedforward-inhibitory components. Paired analysis of simultaneously recorded *St18±* neurons showed similar EPSP amplitudes (EPSP_1_ in GABAzine and CGP 55845: sup, 2.8 mV (1.5, 3.2); deep, 2.7 mV (1.5, 3.6); *n* = 18 pairs, *N* = 3 mice; Wilcoxon signed rank test, *p* = 0.77) and CGP 55845 sensitive responses (measured post-train: sup, −0.1 mV (−4.2, 0.0); deep, −0.4 mV (−4.3, 0.0); *n* = 18 pairs; Wilcoxon test, *p* = 0.42), but stronger GABAzine sensitive responses on *St18−* than *St18+* PNs (PSP_3_ amplitude difference with GABAzine: sup, −1.1 mV (−5.9, −0.3); deep, −0.9 mV (−2.9, 0.1); *n* = 19 pairs; Wilcoxon test, *p* = 0.0017).

Given the non-canonical projection pattern of deep *St18+* PNs within CA3, we sought to characterise the synaptic targets of deep CA3 PNs within the ipsilateral CA3 network. Previous reports have shown that deep CA3 PNs connect preferentially to deep rather than superficial PNs at the local level ^23,61,62^. To understand the contribution of deep *St18+* PNs to the CA3 associational network on a longer-distance range, we performed optogenetic-assisted circuit-mapping in CA3, virally transducing dorsal *St18+* CA3 PNs with ChIEF opsin ^63^, followed by multicellular patch-clamp recording in horizontal slices of mid-ventral hippocampus. We observed short latency light-induced postsynaptic potentials (PSPs) on simultaneously recorded pairs of *St18+* and *St18−* CA3 PNs (**Fig. 4f**). By sequential application of first GABA_A_ and then GABA_B_ receptor antagonists (GABAzine and CGP 55845 respectively), we could isolate excitatory PSPs, which occurred with equivalent amplitudes on both CA3 PN types (**Fig. 4g**; *p* = 0.77). Therefore deep *St18+* PNs provide long-range excitatory recurrent input to both deep and superficial CA3 PNs within the ipsilateral hemisphere. While recording, we observed marked pharmacologically-induced changes to PSP responses. Quantifying these effects, we could confirm variable, but substantial GABAzine and CGP 55845-sensitive responses induced by optogenetic stimulation, consistent with feedforward-inhibitory transmission resulting from deep *St18+* PN activation (**Fig. 4g**). CGP 55845-sensitive responses were similar onto both CA3 PN types, but notably, GABAzine-sensitive feedforward inhibition was approximately twice the magnitude onto superficial *St18−* PNs compared to deep *St18+* PNs (mean: sup, −2.5 mV; deep, −1.3 mV), suggesting preferential inhibitory gating of the superficial layer (**Fig. 4g**). Thus, deep PNs act on their primary output target, the ipsilateral CA3 network, through both excitatory recurrent synapses and preferential feedforward inhibition of the superficial CA3 PN layer.

### Conservation of deep-superficial identity in humans

The striking genetic, functional, and connective differences between CA3 PN types raise the question of whether this architecture represents an evolutionarily conserved principle of hippocampal organisation. To explore conservation beyond mice, we reanalysed existing single-nucleus (sn)RNA-seq datasets from pig and human hippocampus ^64,65^. Leiden clustering of CA2-3 PNs ^64^ from pigs revealed three stable clusters: one expressing canonical CA2 markers (*Rgs14*, *Amigo2*), and two putative CA3 PN subtypes, which clearly segregated into *St18+* and *St18−* populations (**Fig. 5a**). Differentially expressed genes in the pig *St18+* subtype overlapped with those of mice (*St18*, *Efna5*, *Pcdh17*), demonstrating molecular conservation of CA3 PN subtypes across species (**Fig. 5a**).

**Figure 5:**
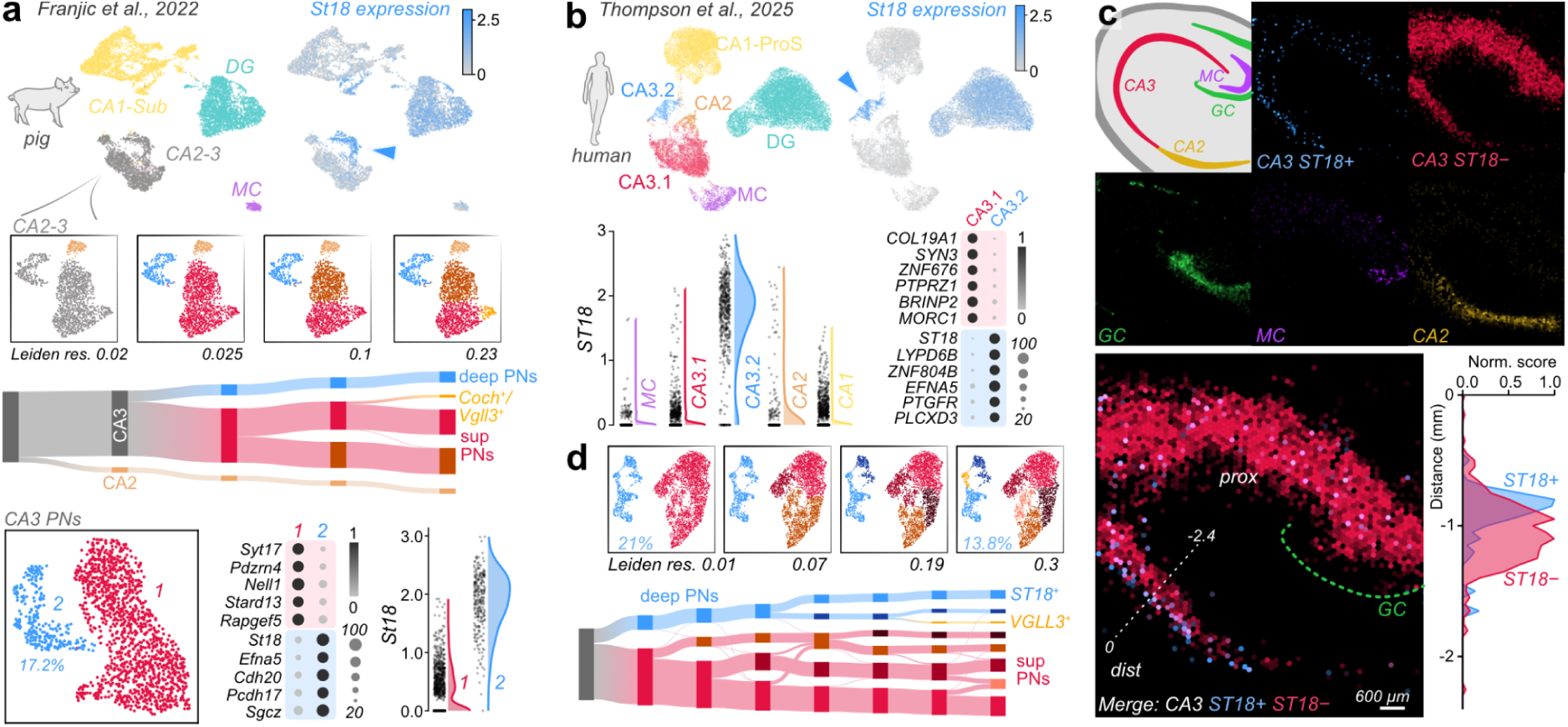
CA3 PN subtypes are molecularly conserved from rodents to humans. **a,** Re-analysis of pig hippocampal snRNA-seq data from Franjic et al. 2022 reveals conserved *St18* expression in a subset of CA3 PNs. Leiden clustering of ‘CA2-3’ at increasing resolutions resolved three stable populations: CA2 PNs (*Rgs14+, Amigo2+*), and two putative CA3 PN subtypes, corresponding to superficial and deep subpopulations. Subclustering of CA3 PNs identified two transcriptomically distinct groups, with the deep cluster selectively expressing conserved markers *St18, Efna5* and *Pcdh17* (dot plot). Violin plot confirms molecular conservation of *St18+* and *St18−* CA3 PNs in the pig hippocampus (units: ln(CP10k+1)). **b,** Re-analysis of human hippocampal snRNA-seq data from Thompson et al. 2025 reveals specific *ST18* expression in the ‘CA3.2’ subtype, one of the two CA3 PN clusters identified in this dataset. UMAP and violin plot (units: ln(CP10k+1)) confirm selective *ST18* enrichment in ‘CA3.2’ relative to ‘CA3.1’, as well as to CA2, CA1, and mossy cells (MC). Differential gene expression analysis further supports the identity of CA3.2 as the deep CA3 PN population, based on conserved expression of deep PN markers including *ST18*, *EFNA5*, and *PLCXD3*. **c,** Spatial transcriptomic data from healthy human tissue (replotted from ^65^) shows conserved deep-superficial organisation of *ST18+* and *ST18−* CA3 PN subtypes. NMF signatures for granule cells (GC), mossy cells (MC), CA2, and *ST18*− CA3 PNs show expected spatial distributions; *ST18*+ CA3 PNs form a thin band in deep, distal CA3. Expression along the indicated line is plotted. **d,** Re-clustering of CA3 PNs from the human dataset confirms early segregation into deep and superficial cell populations. A small *ST18−* subpopulation co-clustering with the deep cells and expressing conserved deep PN specific genes suggests further heterogeneity within human deep CA3 PNs.

To extend this analysis to humans, we used spatially-informed hippocampal snRNA-seq data ^65^. In this dataset, while CA1 and CA2 PNs are represented as individual populations, CA3 PNs are presented as two subtypes: CA3.1 and CA3.2 (**Fig. 5b**). Plotting *ST18* expression on both UMAP (Uniform Manifold Approximation and Projection) representations and across CA neuron clusters demonstrated clear and specific labelling of the CA3.2 subtype, directly recapitulating the CA3 PN subtypes found in mice (**Fig. 5b**). Mapping these subtypes onto spatial transcriptomic data using their distinct non-negative matrix factorization (NMF) signatures showed deep and superficial laminar distributions of *ST18+* and *ST18−* subtypes in distal CA3 (**Fig. 5c**). Therefore, deep *ST18+* and superficial *ST18−* PNs form a conserved two-layer architecture in CA3 in the human hippocampus. For completeness, we extracted and reclustered all CA3 PNs from this dataset, which confirmed clear segregation of *ST18+* and *ST18−* PNs between the primary subgroups (**Fig. 5d**). A small subgroup of *ST18−* neurons expressing conserved deep PN marker genes (*EFNA5*, *PCDH17*) co-clustered with deep *ST18+* PNs in this analysis, suggesting the possibility of additional heterogeneity within deep PNs in humans that requires future exploration (**Fig. 5d**).

Given the genetic conservation of deep and superficial CA3 PNs in humans, we tested whether we could identify these cell types functionally. Temporal lobe epilepsy resections provide a unique opportunity to record from live human hippocampal tissue, and we exploited this resource using non-sclerotic samples, where hippocampal architecture is preserved ^66–68^. We performed multicellular patch-clamp recordings from distal CA3, simultaneously recording PNs spanning the deep-superficial axis (**Fig. 6a**). We observed higher burst firing, the functional hallmark of deep PNs in mice ^52,17,21,23^, in a subpopulation of human CA3 neurons whose somata were confined to deep positions in the pyramidal cell layer (**Fig. 6b**). We used this metric alone to assign PNs to putative deep or superficial subclasses for further analysis (**Fig. 6a-b**; **Supplementary Fig. 5**).

**Figure 6:**
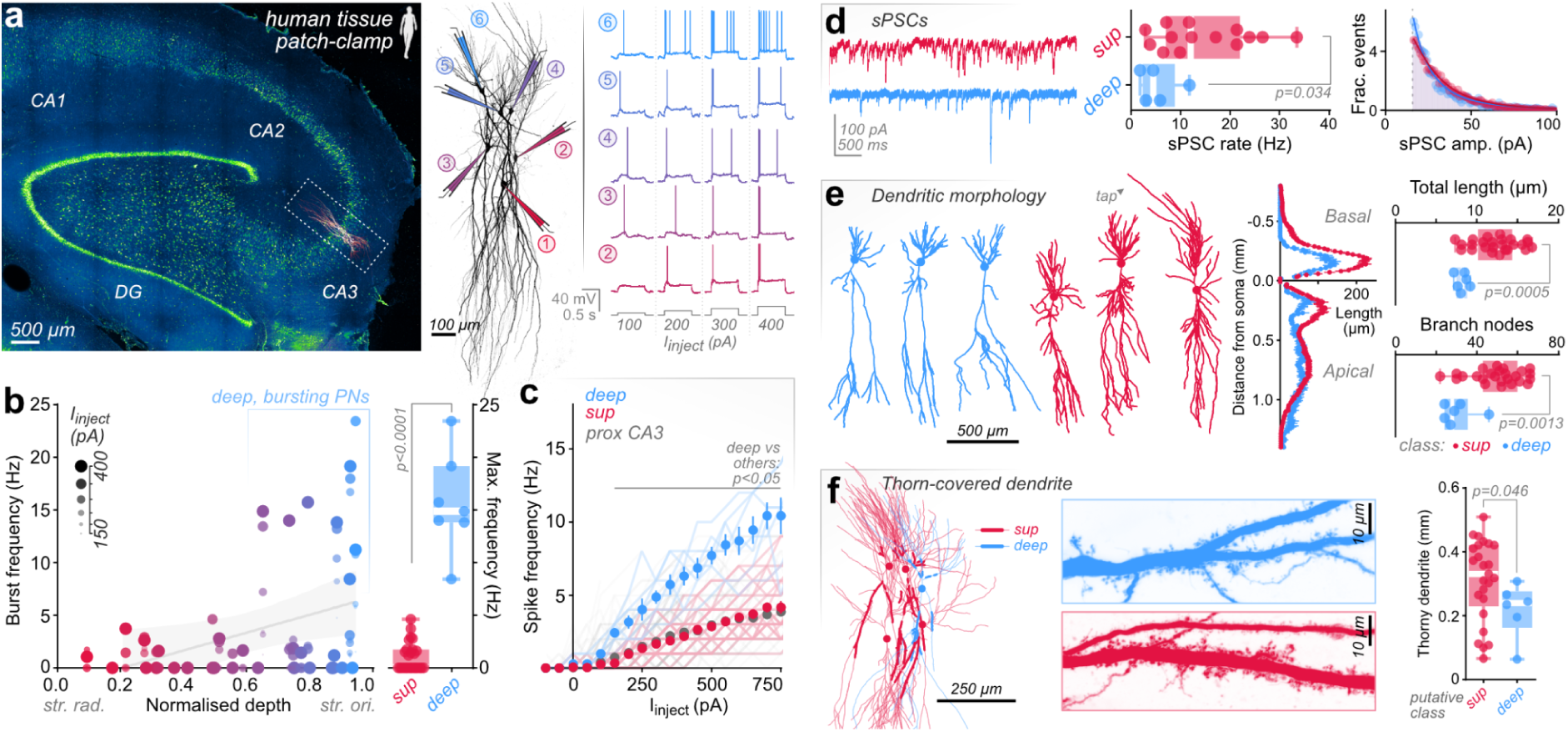
CA3 PN subtypes show conserved functional properties in humans. **a**, Example patch-clamp recording from human hippocampal CA3, showing recording configuration spanning the deep (blue) to superficial (red) axis, and representative firing profiles upon current injection. **b**, Burst frequency (first three spikes) plotted against normalised somatic depth across current injections (circle size) reveals a subpopulation of burst-firing PNs restricted to deep in the pyramidal cell layer (partially boxed). Linear regression of 400 pA current injection (grey) was significantly positive (R^2^ = 0.145, *p* = 0.029; *n* = 33 neurons, 6 recordings, 2 patients). Maximum firing frequency across all current injections (right) was markedly higher in putative deep than superficial populations (median (IQR): sup, 0.0 Hz (0.0, 1.7), *n* = 26; deep, 14.9 Hz (13.8, 19.2), *n* = 7; Mann-Whitney, *p* < 0.0001). **c**, Superficial neuron spike frequency-current relationships were comparable to previously recorded proximal human CA3 PNs ^68^, while deep PNs showed steeper F-I curves (mixed-effects analysis, *p* < 0.0001; Tukey’s multiple comparisons test *p* < 0.05 for deep vs sup and deep vs prox CA3 from 150 pA injection). **d**, Spontaneous postsynaptic current (sPSC) rates were much higher on sup than deep PNs, echoing mouse PN types (sPSC rate: sup, 12.0 Hz (7.0, 22.0), *n* = 14; deep, 4.5 Hz (2.4, 8.9), *n* = 5; *p* = 0.033). Relative frequency histograms of sPSC amplitude show subtle skewing to lower amplitudes on deep PNs (lognormal fit, extra sum-of-squares F-test, *p* < 0.0001; 1 pA bin width with detection limit (grey) calculated as 15.7 pA ^69^). **e**, Reconstructed dendritic morphologies of putative subclasses (example with a ‘taproot’ basal dendrite marked ‘tap’) showed reduced dendritic complexity for deep than superficial neurons (total dendritic length, median (IQR): sup, 12.8 mm (10.2, 14.4), *n* = 25; deep, 8.3 mm (7.5, 8.8), *n* = 6; Mann-Whitney, *p* = 0.0005; number of branch nodes per tree: sup, 52 nodes (43, 60); deep, 28 nodes (24, 35.5); *p* = 0.0013), with Sholl analysis showing reduced branching complexity in both basal and apical dendrites close to the soma (10 μm bin-width; symbols depict mean ± SEM). **f**, All human CA3 PNs exhibited thorny excrescences, with the length of thorn-covered dendrite modestly reduced on deep PNs (sup, 331 μm (230, 421), *n* = 25; deep, 240 μm (163, 277), *n* = 6; *p* = 0.046), following total dendritic length.

Putative deep and superficial human CA3 PNs differed in multiple morpho-electric parameters, recapitulating subtype-specific features observed in mice. Deep PNs showed significantly higher firing rates, lower membrane capacitance, lower rheobase current, and a markedly lower rate of spontaneous PSC (sPSC) input compared to superficial PNs (**Fig. 6c-d, Supplementary Fig 5**). Reconstructed morphologies also showed subtype-specific differences, with a lower dendritic length and significantly fewer branch points per neuron for deep than superficial PNs (**Fig. 6e**). Branching was reduced in both basal and apical compartments close to the soma in deep PNs, consistent with a preferential reduction in CA3 recurrent, rather than entorhinal cortical input to these cells (**Fig. 6e**). Within cell classes, none of these features correlated with somatic depth, and restricting our analysis to deep and superficial PNs with equivalent somatic depth showed significant differences in all subtype-specific features (**Supplementary Fig. 5f**). Together, these findings support discrete cell type identity of deep, bursting PNs, rather than graded laminar properties amongst the human CA3 neuron population.

Despite PN subtype conservation, we observed several distinct features in human CA3. Somatic density is far sparser in humans, with less distinct lamination than the mouse hippocampus. As a result, the hallmark ‘long-shank’ of deep CA3 PNs was not observed in humans (**Supplementary Fig. 5e**). Taproot basal dendrites, occurring in humans but not in mice ^68^, were observed only on superficial PNs, adding further to their dendritic complexity (**Supplementary Fig. 5e**). Lastly, while the total dendritic length bearing thorns, receiving apparent DG input, was lower in human deep PNs (**Fig. 6f**), this was proportional to their reduced overall dendritic length (**Supplementary Fig. 5e**), and completely ‘athorny’ PNs were not observed. Reduced, but not absent, mossy fibre input is thus a conserved circuit feature of deep CA3 PNs across species.

Our findings demonstrate that CA3 is formed of two distinct PN subtypes, conserved across species, forming a fundamental circuit element in the mammalian hippocampus. By understanding CA3 PN heterogeneity in this way, we can add circuit context to an intriguing initial observation by Thompson et al. ^65^. Of all hippocampal cell types, CA3.2 cells alone showed significant association with intelligence and educational attainment in human genomic analysis. Our data identify these cells as deep *ST18+* cells; molecularly and functionally distinct CA3 PNs forming a non-canonical arrangement in the autoassociative processor. This element may therefore be a circuit feature that generates differences in human cognitive ability.

## DISCUSSION

CA3 PNs provide the majority of synapses to the hippocampal system, a projection architecture that has inspired current theories of associational memory ^10,11,13^. In the classical model, CA3 forms one broad recurrent network across hemispheres, capable of associating and storing diverse input, and projecting downstream to CA1. Our data prompt a revision of this map.

CA3 is formed of two genetically distinct PN subtypes conserved across mammals: superficial *St18−* (type I) PNs, and deep *St18+* (type II) PNs ^26^. More abundant superficial *St18−* PNs form the textbook autoassociative network and the second step in the canonical trisynaptic circuit between DG and CA1 ^12^. Deep *St18+* PNs receive less DG input, provide less feedforward projection to CA1, and almost none to the contralateral hippocampus, functioning in some ways more like a CA3-focussed excitatory interneuron than a classical projection neuron. These cells comprise a previously unrecognised circuit module within the associative network, conferring computational capabilities absent from the existing CA3 model ^12^. Deep PNs receive input from the superficial PN autoassociative network, entorhinal cortex in *stratum lacunosum moleculare* ^17,23^, and reduced, but not absent input from the DG. Within the recurrent network, deep PNs preferentially receive synaptic input from the superficial layer at the local level ^23,62,61^, forming their own recurrent network that projects to both layers at longer distances within each hemisphere. Inhibition is also specifically wired in this network. Deep and superficial subnetworks receive independent perisomatic inhibition, allowing layer-specific functional control ^18,23^, while deep PN activation drives particularly strong feedforward inhibition onto the superficial PN layer.

This CA3-focussed circuit architecture is consistent with a primary role for the deep PN network as a regulator of the powerful superficial associational processor. Its network positioning could curb overexcitation in the superficial recurrent network through feedforward inhibition, while its excitatory connectivity could broadcast completed superficial PN ensembles (the result of CA3 pattern completion ^14,9^), forming ‘association of associations’, or propagation of ensemble sequences ^13^. Both CA3 PN subtypes are strongly modulated by cholinergic input, which increases superficial and decreases deep PN activity^17,22^. This property alone, or combined with layer-specific inhibition, allows brain-state dependent engagement of the deep PN network. Deep PNs are likely to be disengaged during exploration and memory acquisition, and active during periods of memory reactivation, where they are thought to promote sharp-wave ripples ^17^. In this model, ensemble learning could occur in the superficial associational network without deep PN interference, yet memory reactivation, with its characteristic temporally compressed sequence firing, would be driven by burst-firing of deep *St18+* PNs. Segregation of ‘read’ and ‘write’ modes in hippocampal information processing has been proposed ^70,71^. The deep PN circuit offers a potential substrate to achieve this separation.

Another surprising feature of deep PN wiring is their axonal pattern, projecting primarily at the border between *stratum radiatum* and *stratum lacunosum moleculare*. Recent work has identified a distinct electrophysiological signal at this location, which may be the signature of deep PN activity ^60^. The *radiatum*-*lacunosum* border is highly enriched in interneuron somata ^59^, a likely mechanism for the strong feedforward inhibition of superficial PNs that we observe following optogenetic activation of deep PNs. Situated between more soma-proximal recurrent CA3 inputs and the distal entorhinal cortical inputs, deep PN axons are positioned to control coupling between the internal hippocampal associative network and external EC-driven input on these actively integrating apical dendrites ^20,72^. This arrangement is reminiscent of the dendritic gating mechanisms proposed in both CA1 ^73^ and neocortical PNs ^74^. State-dependent modulation of deep PN activity could therefore dynamically regulate the extent to which external sensory context influences ongoing associative computations.

The largely unilateral wiring of deep PNs is particularly intriguing. This finding is independently supported by whole brain projectome data ^19^, in which CA3 PNs project bi-and unilaterally in a 75:25 ratio — approximating the proportions of *St18+* and *St18−* PNs. This wiring architecture provides a circuit mechanism for hemisphere-specific functions to be imposed on an otherwise bilaterally acting processor. The superficial CA3 recurrent network is built across hemispheres, allowing powerful association of potentially all hippocampal inputs ^6,12^, yet hemisphere-specific functions in humans have been widely reported ^75,76^. Even in mice, left and right CA3 projections make functionally distinct contributions to spatial memory formation ^77,78^. Regulation of the CA3 recurrent processor by distinct deep PN circuits in each hemisphere may therefore provide a circuit basis for hippocampal functional lateralisation.

Our findings contribute to a growing picture of parallel computation in hippocampal circuits ^38,79,80^. Despite PN heterogeneity being much better studied in CA1 ^34–38,40^, our data demonstrate a far greater distinction between deep and superficial subtypes in CA3. In CA1, PNs form a genetic continuum with deep CA1 PNs in one corner of a complex landscape, rather than forming discrete subtypes as we observe in CA3. This idea is supported by functional measurements in human CA1, where morpho-electric diversity lacks clear subtype distinctions ^67^. Projection patterns of deep and superficial CA1 PNs are also much more similar than the highly non-canonical projections we observe from deep CA3 PNs. CA3 PNs therefore appear in many ways similar to layer 5 of the neocortex, where extratelencephalic (ET) and intratelencephalic (IT) PNs occupy the same layer, yet have highly distinct genetics, projections, and circuit roles ^81^. Deep *St18+* and superficial *St18−* CA3 PN subtypes fulfil all four criteria for cell type identity — morphological, electrical, transcriptomic, and connective (METC-types) — with the two-layer architecture extending the full dorso-ventral extent of the hippocampus. We therefore propose that the deep-superficial axis is the primary organising principle of CA3, with dorso-ventral functional differences ^31^ likely reflecting differential input to a conserved circuit architecture. Proximal CA3 may represent a partial exception, with fewer *St18+* PNs, a lack of EC input ^8^, and superficial PNs with distinct genetic and functional properties ^23,27,30^.

Genetic identification of CA3 PN subtypes provided the molecular lens through which to identify a previously unknown circuit feature, demonstrating the importance of understanding genetic cell type identity. This perspective also offers an opportunity to understand the molecular basis for PN subtype differences. The deep PN marker gene *St18* is a transcription factor determining deep layer neuronal identity even in salamanders ^82^, and it induces ‘projection-like’ rather than collateralising axon patterns in mouse interneurons ^83^. It is therefore a prime candidate to mechanistically determine the distinct projection patterns of CA3 PN subtypes, and conservation of *St18* across mice, pigs, and humans also suggests an important role for the gene in circuit function. Multiple other genes that are differentially expressed between CA3 PN subtypes have circuit sculpting roles (*Efna5*, *Efnb3*, *Epha3*). *Efna5*, a conserved deep-specific marker, is a candidate to generate intrahemispheric projections, given its role in generating commissural fibre projections ^84^ and axonal pruning ^85^. *Cdh13* controls differential inhibitory wiring between layer 5 PN subtypes in the neocortex ^86^, suggesting a parallel role in CA3’s layer-specific inhibitory topology. Further synaptic (*Cacng2*, *Cacng3*, *Olfm1*, *Gabrg3*, *Homer3*) and neuromodulatory receptor genes (*Htr2c*) offer PN subtypes the potential for cell type-specific plasticity mechanisms that can now be explored. *St18*-Cre animals provide a key tool to understand these potential functions.

Together, by identifying the primary axis of CA3 PN heterogeneity, and viewing the circuit through this lens, we reveal a two-layer architecture that fundamentally revises our understanding of the hippocampal associational circuitry. The exact mechanistic role of the deep *St18+* CA3 PN network now stands as a major open question. Given that amongst all hippocampal neurons, deep CA3 PNs are uniquely enriched in genetic variants associated with cognitive abilities, this intriguing circuit element may be a key site through which genetic variation shapes human learning and memory capacity.

## METHODS

### Animals

All procedures were performed in strict accordance with institutional, national, and European guidelines for animal experimentation, approved by the Bundesministerium für Bildung, Wissenschaft und Forschung of Austria (Project ‘Connectomics’: GZ 2020-0.648.587 and 2024-0.904.911).

*St18*-3’iCre mice (codon-improved Cre ^50^, referred to henceforth as *St18*-Cre) were generated in house, as detailed below. Ai9 reporter mice (RRID:IMSR_JAX:007905) were crossed with *St18*-Cre to generate heterozygous *St18*-Cre:Ai9 animals. *Ngn2*-CreER(hem):Ai14(het/wt) (MGI:2652037 ^57^; RRID:IMSR_JAX:007914) male and female mice were obtained from Rosa Cossart (INSERM, Marseille, FR). Both *Ngn2*-CreER(hem):Ai14(hom) and *St18*-Cre(het):Ai9(hom) male mice were crossed with wild-type CD1 (Charles River; RRID:IMSR_CRL:022) female mice to generate mice with equivalent genotypes, but on a mixed C57BL/6J, CD-1 background, which was more resilient to tamoxifen and BrdU injections during pregnancy. *St18*-Cre mice with C57BL/6J background (RRID:IMSR_JAX:000664) were used throughout the study, while all experiments requiring tamoxifen or BrdU injections were performed on this mixed C57BL/6J:CD-1 background. Experiments used *St18*-Cre(het), *St18*-Cre(het):Ai9(hom/het) and *Ngn2*-CreER(hem):Ai14(het) genotypes throughout.

All animals were housed with ad libitum access to food and water, with constant temperature and humidity of 22°C and 50–60%, respectively, under a 12-hour light-dark cycle, and used for experiments during the light phase.

### Generation of St18-Cre mice

*St18*-Cre mice were generated by the ISTA Transgenic Unit on a C57BL/6J genetic background by 2-Cell Homologous Recombination-CRISPR ^87^. Zygotes were microinjected with CRISPR mix consisting of Cas9 mRNA, single guide RNA (sgRNA) targeting the 3’ end of the *St18* protein encoding region and a custom designed repair template. The repair template consisted of 1 kb homology arms on either side of the sgRNA binding site (TGTGTAGAACGACAGTATGC<u>AGG</u>) with an inserted sequence encoding a self-cleaving T2A peptide followed by iCre immediately before the endogenous *St18* stop codon.

Three to four week-old C57BL/6J donor females were hormone primed by intraperitoneal injection of 5 IU (international units) of pregnant mare serum gonadotropin (PMSG) to induce superovulation. 5 IU of human chorionic gonadotropin (hCG) was administered 46–48 h after PMSG treatment, followed by one to one mating with stud males. Zygotes were isolated from the oviduct of mated females at 0.5 days post coitum. M2 medium with hyaluronidase (MR-051-F, Sigma) was applied briefly to dissociate cumulus cells, before culturing zygotes in small drops of KSOM embryo culture medium (MR-121-D, Sigma) under mineral oil (M-5310, Sigma) at 37°C with 5% CO_2_, before microinjection with the CRISPR mix detailed above. Genetic modification of F0 founder mice was confirmed by PCR amplification using primers spanning homology arms, and F1 mice were generated by backcrossing founder mice with C57BL/6J animals. F1 mice were genotyped by PCR, separately amplifying both T2A-iCre and homology arms. Positive PCR products were Sanger sequenced to confirm precise targeting.

### Neuronal birthdating using Ngn2-CreER and BrdU

To induce CreER activity, tamoxifen (10 mg⋅mL^−1^ in corn oil; 2 mg/30 g body weight) was administered intraperitoneally to wild-type CD1 pregnant dams crossed with *Ngn2*-CreER(hem):Ai14(hom) males, at E12.5, E14.5, and E16.5 post vaginal plug (E0.5). Dams were allowed to produce litters, and offspring were used for experiments at postnatal day (P) 20–65.

For BrdU labelling, *St18*-Cre(het):Ai9(hom) males were crossed with wild-type CD1 females, and pregnant dams received a single intraperitoneal injection of BrdU (10 mg⋅mL^−1^ in saline; 3 mg/30 g body weight). Again, dams were left to litter down, and offspring were used for experiments at P43–61.

### Viral vectors

For projection labelling, AAV-CAG-DIO-tdTomato (Addgene plasmid #28306, courtesy of Edward Boyden) and AAV-pCaMKII-SEO-EGFP (derived from Addgene plasmid #172361) constructs were used, while AAV-DIO-EF1a-ChIEF-EYFP was used for optogenetic experiments. All viruses were packaged into capsids of AAV-DJ serotype ^88^. AAV payload, AAV-DJ Rep/Cap, and pHelper (Cell Biolabs) plasmids were cotransfected into Lenti-X 293T cells (Takara Bio, Cat. #632180; RRID: CVCL_4401) at 80–90% confluency. 72 h after transfection, cells were harvested, washed with phosphate-buffered saline (PBS), and lysed in AAV lysis buffer (150 mM NaCl, 50 mM Tris-HCl, pH 8.0, with 2 mM MgCl_2_, and 0.5% sodium deoxycholate). Samples were subjected to three freeze-thaw cycles before Benzonase treatment at 37°C for 45 min (50 U⋅ml^−1^), centrifugation at 2000 ×g for 30 min at 4°C, and purified on a discontinuous iodixanol gradient (17, 25, 40, and 60% fractions) by ultracentrifugation at 350,200 ×g for 90 min. The AAV-containing 40% fraction was collected and concentrated using 100-kDa molecular weight cutoff centrifugal filters (Sartorius, Vivaspin 20), before titration using qPCR, and dilution in 0.9% saline (PARI, 077G0020) for injection at 1–5×10^12^ GC⋅ml^−1^.

### Stereotactic injections

For *in vivo* delivery of viral vectors, animals were anaesthetised with isoflurane, administered subcutaneous buprenorphine 0.1 mg kg^−1^ (Buprenidine, Dechra Veterinary Products, 836699), and placed in a custom stereotactic frame on a heat pad with continuous delivery of 2–4% isoflurane (1 L·min^−1^ O_2_ flow). When toe pinch reflexes were no longer observed, a skin incision was made to expose the skull above the scalp. Bregma was used as the reference coordinate for anterior-posterior (A-P) and medio-lateral (M-L) injection coordinates, and a small hole was drilled through the skull at this location, while the surface of the brain was used as the reference coordinate for the dorso-ventral (D-V) axis. In all experiments, dorsal CA3 was targeted with the coordinates −1.9 mm A-P/ ±2.5 mm M-L/ −2 mm D-V. Viruses were delivered using a Nanoject III system (Drummond Scientific) fitted with a borosilicate glass pipette (Drummond, #3-000-203-G/X) backfilled with mineral oil (Sigma-Aldrich; M5310), mounted on a Luigs & Neumann Mini-25 manipulator. After injection, the pipette was left in position for 5 min before slowly retracting, to allow the virus to diffuse into the tissue and to prevent solution retraction up the pipette tract. After injection, the scalp wound was glued together (Vetbond, 3M) and mice were returned to their home cage to recover.

For analysis of projections, *St18*-Cre mice aged between 4 and 14 weeks of both sexes were injected unilaterally with a mixture of AAV-CAG-DIO-tdTomato and AAV-CaMKIIa-SEO-EGFP. AAVs were diluted 1:5 from stocks in 0.9% saline (PARI, 077G0020) and mixed 1:1, immediately before injection. 400 nL viral mixture was injected at the desired coordinates, at a speed of 2–4 nL·s^−1^. Injections were performed in both left and right hemispheres, and as no difference between hemispheres was observed, data were pooled.

For optogenetic stimulation experiments, *St18*-Cre:Ai9 mice aged P24–25 were injected in the left dorsal CA3 with AAV-DIO-EF1a-ChIEF-P2A-EYFP in an equivalent manner, using 200 nL of diluted viral solution. Animals were used to produce acute slices as detailed below, with recordings performed 10–19 days after injection.

### Histological analysis of viral, genetic, and birthdated neurons

Animals were euthanised by transcardial perfusion first with phosphate buffer (PB; 0.1 M NaH_2_PO_4_ and 0.1 M Na_2_HPO_4_ (Merck) titrated to pH 7.35), and then with 4% paraformaldehyde (PFA) in PB (v/v from 16% PFA; TAAB Laboratories Equipment) under terminal anaesthesia. For anaesthesia, an intraperitoneal injection was administered of either MMF (0.3 mg kg^−1^ medetomidine, Dormitor, Pfizer; 8 mg kg^−1^ midazolam, Dormicum, Roche; and 0.01 mg kg^−1^ fentanyl, Piramal Critical Care) or ketamine/xylazine/metamizol (100 mg kg^−1^ Ketamidor, VetVita, 8-01141; 10 mg kg^−1^ Rompun, Elanco Animal Health GmbH, 14840; 200 mg kg^−1^ Novalgin, 500 mg⋅ml^−1^, Sanofi, 3.191; in 0.9% NaCl). Brains were harvested intact, and postfixed by overnight immersion in 4% PFA/PB at 4°C. After 24 h, the fixation reaction was terminated by washing with 0.1 M PB, and brains were sliced in PB to 50–100 μm thick sections using a Leica VT1200 vibratome. Additionally, some BrdU-labelled sections were embedded in 2% agarose (0.5 g of agarose, Sigma A9539, in 25 mL of water; microwaved until dissolved, and cooled down before transfer) before sectioning. Coronal or horizontal cutting planes were typically used for analysis of dorsal and ventral hippocampus, respectively.

Immunohistochemical analysis was performed where appropriate on sectioned tissue. Sections were washed in PB, blocked in 10% normal goat serum (NGS; Biozol ENG9010-10) in PB, and permeabilized by incubation with a solution containing 5% NGS and 0.4% Triton X-100 in 0.1 M PB for 1 h at room temperature (RT). Primary antibodies were then added in blocking/permeabilizing solution overnight at RT with gentle shaking. Slices were washed (3 × 10 min in PB) before addition of secondary antibodies in blocking/permeabilizing solution, incubated overnight at RT with gentle shaking. Slices were washed again (3 × 10 min in PB) before mounting in Mowiol (Mowiol 4–88, Carl Roth, 713.2) or Prolong Gold (Invitrogen, P36930) between a glass slide (Assistent, Karl Hecht ref. 42406020) and a 1.5 thickness glass coverslip (VWR 631-0147).

For analysis of *St18* and *Ngn2* cell distribution, guinea pig anti-NeuN (Sigma-Aldrich, ABN90; RRID:AB_11205592) primary antibody, and AF647-conjugated goat anti-guinea pig IgG (Thermo Fisher Scientific, A-21450; RRID:AB_2535867) secondary antibodies were used, both diluted 1:500. Half hemispheres were sectioned using coronal and horizontal cutting planes. For CA3, CA1 and neocortex distributions, slices were imaged using the ANDOR Dragonfly microscope (Oxford Instruments) equipped with a Zyla 4.2 Megapixel sCMOS camera (2048 x 2048 pixels). Images of the hippocampus and adjacent neocortex were taken using a 20x water-immersion objective (Nikon MRD77200, CFI P-Apochromat 20x, NA 0.95) with 1 μm z-steps. For the radial distribution analysis of *St18* CA3 PNs in **Fig. 1d**, higher magnification images were taken using a 40x water-immersion objective (Nikon MRD77410, Apochromat LWD λS 40x, NA 1.15) with 0.5μm steps. Pinhole discs with 25 or 40 μm hole diameters were used for 20x or 40x objective images, respectively. Tiles were stitched using Imaris Stitcher software (Oxford Instruments). Stitched image stacks were converted to TIFF format using FIJI ^89^. *Stratum lucidum* and *stratum oriens* borders of CA3 and CA1, as well as white matter and layer I borders within the neocortex, were then delineated using Neutube analysis software ^90^. Neutube was also used to mark *St18*+, NeuN+ and *Ngn2*+ cells for distribution analysis, generating SWC files that were then subsequently analyzed using custom MATLAB scripts.

For BrdU cell distribution and co-localization with *St18* analysis, coronal and horizontal-cut slices were first incubated in 1 M HCl at 45°C for 30 minutes, before washing twice with 0.1 M PB. They were then incubated for 10 minutes in borate buffer (100 mM boric acid, 75 mM NaCl, and 25 mM sodium tetraborate), before being washed in 0.1 M PB with 0.4% Triton, and subsequently blocked and permeabilized with primary antibodies as described above. The primary antibodies used were rat anti-BrdU (1:1000, Abcam ab6326, RRID:AB_305426) and rabbit anti-NeuN (1:500, Thermo Fisher Scientific PA5-37407, RRID:AB_2554049), with secondary AF488-conjugated chicken anti-rat (Thermo Fisher Scientific, A-21470; RRID:AB_2535873) and AF647-conjugated goat anti-rabbit (Thermo Fisher Scientific, A-21244; RRID:AB_2535812) both diluted 1:500. Images of the hippocampus and neocortex were acquired with a 20x water-immersion objective, stitched, and analyzed in Neutube as described above. Additionally, the number of BrdU+ and *St18*+ co-localized cells was quantified in FIJI.

For analysis of projections, chicken anti-GFP (Abcam ab13970; RRID:AB_300798) and mouse anti-RFP (Thermo Fisher Scientific, MA5-15257; RRID:AB_10999796) primary antibodies, and AF488-conjugated goat anti-chicken IgY (Thermo Fisher Scientific, A-11039; RRID:AB_2534096) and AF568-conjugated goat anti-mouse IgG1 (Thermo Fisher Scientific, A-21124; RRID AB_2535766) secondary antibodies were used (all diluted 1:500 from stock). Ipsi- and contralateral projections were compared from fluorescence line profiles in CA3 both in caudal coronal sections (approximately −2.7 mm from Bregma) and horizontal sections, ensuring that no dendrites of labelled cells were included. Line profiles from ipsi- and contralateral CA3 of the same slice and image were spatially binned (12 bins in *stratum oriens*, 20 in *stratum radiatum*), background subtracted, and normalised to the peak fluorescence value, and the ratio of ipsi- to contralateral peak fluorescence was taken as the projection ratio. CA1 vs CA3 projections were compared in horizontal sections only, to avoid dendritic fluorescence at the injection site, and to compare profiles in planes perpendicular to the long axis of the hippocampus. Fluorescence line profiles from CA3 and CA1 of the same slices were acquired and normalised in the same manner as ipsi-contra comparisons, where the peak binned fluorescence was used to compute the ratio of CA1:CA3 innervation. Presented data includes analysis of axons from both DIO-tdTomato/SEO-EGFP injections, and from acute slices with DIO-ChIEF/SEO-ChIEF expression after optogenetic recording. Paired data present comparisons of PN subtype projections in the same animal and slice.

### Human patient tissue samples

Human tissue samples were obtained with informed patient consent from two individuals, with right hippocampal tissue resected during treatment of temporal lobe epilepsy, and a low grade glioma (tumour-related epilepsy). This work was approved by the Ethics Committee of the Medical University Vienna (MUW) (EK Nr: 2271/2021). No evidence of hippocampal sclerosis was observed during pre-surgical MRI, or post-surgical neuropathological analysis, and therefore samples were considered ‘non-sclerotic’ ^66,68^. Both patients were female, aged 25 and 47 years, respectively.

### Human hippocampal slice preparation

The hippocampus and parahippocampal gyrus (PHG) were surgically harvested by anteromesial temporal lobe resection (AMTLR), avoiding any mechanical or hypoxic damage to the tissue. The hippocampus/PHG was prepared such that the vascular supply was disconnected just before removal, so that an “en bloc” specimen was transferred immediately to the transport solution. A substantial portion of resected tissue was sent to the neuropathological laboratory (Division of Neuropathology and Neurochemistry, Department of Neurology, Medical University of Vienna, MUW) for complete histological assessment, neuropathological diagnosis, and sub-classification according to international league against epilepsy (ILAE) sclerosis standards ^91^.

Tissue blocks were immediately submerged in ice-cold high-sucrose artificial cerebrospinal fluid (aCSF, containing 64 mM NaCl, 25 mM NaHCO_3_, 2.5 mM KCl, 1.25 mM NaH_2_PO_4_, 10 mM D-glucose, 120 mM sucrose, 7 mM MgCl_2_, and 0.5 mM CaCl_2_; osmolarity ∼334 mOsm) equilibrated by bubbling with 95% O_2_ and 5% CO_2_ gas mixture (carbogen). Samples were transported from the operating theatre to the laboratory on ice with continuous carbogen bubbling, taking approximately 45 min. Upon arrival, tissue blocks were trimmed to remove the PHG and form a glueing face perpendicular to the longitudinal axis of the hippocampus (parallel to dendritic arbours). Tissue blocks were glued to the specimen plate of a Leica VT1200 vibratome (Leica Microsystems) with liquid superglue (UHU), and submerged in a partially frozen ‘slush’ of ice-cold high-sucrose aCSF with supplemental carbogen bubbling. 350-μm-thick acute slices were cut (blade advance velocity 0.04 mm⋅s^−1^, oscillation amplitude 1.2 mm), before recovery at 35°C in high-sucrose aCSF for 30–45 min. Slices were recovered and recorded with their cut face downwards. After recovery, slices were maintained in bubbled high-sucrose aCSF at RT (20–22°C) until recording. Slices were recorded within 12 h of sectioning.

### Mouse hippocampal slice preparation

For electrophysiology wild-type C57BL/6J (RRID:IMSR_JAX:000664), *St18*-Cre:Ai9, or *Ngn2*-CreER:Ai14 mice from both sexes were used at P20−55. For those aged <P30, animals were euthanised by decapitation under isoflurane anaesthesia, while those ≥P30 were euthanised by transcardial perfusion of 15 mL ice cold high-sucrose solution equilibrated with carbogen (see above) under terminal anaesthesia (MMF or ketamine/xylazine/metamizol). Brains were then rapidly removed, and immersed in ice-cold high sucrose solution. A single sagittal cut was made to separate hemispheres, before a single horizontal blocking cut to provide a gluing surface at the dorsal face of the brain. Tissue blocks were glued to the specimen plate of a Leica VT 1200 vibratome using liquid superglue (UHU), before sectioning as per human samples, into 350-μm-thick acute slices, and slice recovery at 35°C (see above). Due to this procedure, slices therefore originate from mid to ventral hippocampus.

### Electrophysiology

Slices were transferred to the recording chamber and continuously perfused under gravity flow with recording aCSF (containing 125 mM NaCl, 25 mM NaHCO_3_, 2.5 mM KCl, 1.25 mM NaH_2_PO_4_, 25 mM D-glucose, 2 mM CaCl_2_, and 1 mM MgCl_2_, osmolarity ∼317 mOsm) continuously bubbled with carbogen. Slices were held in place with a platinum harp with nylon threads to prevent tissue movement during recordings, with strings crossing the fimbria and subiculum (avoiding the hippocampus) for mouse slices, and avoiding the CA3 area for human slices. Patch-clamp recording pipettes were pulled from thick-walled borosilicate glass tubing (Hilgenberg, 2 mm OD, 1 mm ID, 1807542), and filled with intracellular solution (containing 135 mM K gluconate, 20 mM KCl, 0.1 mM EGTA, 2 mM MgCl_2_, 2 mM Na_2_ATP, 0.3 mM NaGTP, and 10 mM HEPES, adjusted to pH 7.28 with KOH; osmolarity ∼302 mOsm, with 0.2% (w/v) biocytin). Micropipettes had open-tip resistances of 2–6 MΩ when filled with internal solution, and were positioned manually using up to eight LN mini 25 micromanipulators (Luigs and Neumann). Neurons were targeted using IR-DIC videomicroscopy based on their soma location in the CA3 pyramidal cell layer, and confirmed to be CA3 PNs based on firing properties and post-hoc morphological analysis. In both mouse and human slices, recordings were made from the distal region of CA3 (closer to CA2). Electrical signals were recorded using four Multiclamp 700B amplifiers (Molecular Devices), low-pass filtered at 6–10 kHz with built-in Bessel filters, and digitized at 20 kHz with Power 1401 data acquisition interfaces (Cambridge Electronic Design). Protocols were generated and applied using Signal 6.0 software (CED). Pipette offset and capacitance were measured and accounted for during all recordings. During current-clamp recordings, pipette capacitance was 70% compensated in all cases, and series resistance compensation was applied as appropriate. All recordings were performed at RT (21°C, range: 20–22°C). Neuronal firing properties and membrane properties were assessed by injection of 1-s hyperpolarizing or depolarizing currents in 50 pA steps immediately after achieving the whole-cell configuration. All voltage-clamp recordings were performed with a holding potential of −70 mV. After recordings, pipettes were slowly retracted from the cell somata to form outside-out patches to ensure retention of intracellular biocytin for post-hoc staining. The quality of patch formation and physical location of recorded cells in the tissue slice was documented for later cell identification in stained tissue.

Recorded traces were analyzed using Stimfit (version v0.15.8 ^92^) or custom Matlab scripts (https://github.com/jakefwatson/patchanalysis). Cells with a membrane potential above −50 mV or requiring current injection to maintain membrane potential (V_m_) below −50 mV were excluded from analysis. Resting membrane potential was measured as the median V_m_ without current injection (0 pA injection step of firing characterization), input resistance was calculated from the end of −100, −50, and +50 pA injections (median voltages between 0.65 and 0.95 s of 1-s pulses). Rheobase current was the required injection for first spike generation during 50-pA step injections. Maximum bursting frequency was defined as the highest instantaneous frequency of the first two spikes upon current injection at either 350 or 400 pA steps for analysis of mouse responses. Due to the more temporally restricted spiking pattern of human CA3 PNs ^68^, bursting frequency was quantified from the first three spikes in human tissue. Apparent membrane time constant was measured as a monoexponential fit of the ‘off step’ of hyperpolarizing current injections, and used to calculate cell capacitance. The AP threshold criterion was set as dV/dt = 20 V·s^−1^, and half-duration was measured from resting membrane potential. sPSCs were recorded for 400 s per cell, monitoring voltage-clamp conditions with a test pulse every 20 s. Cells that had a series resistance over 20 MΩ were discarded, and the series resistance of included cells was no different between conditions (human CA3 PNs: sup, 14.8 ± 3.6 MΩ, *n* = 14; deep, 14.2 ± 5.1 MΩ, *n* = 5; Mann-Whitney, *p* = 0.89). Events were detected using Eventer ^93,94^ (fast Fourier transform (FFT)-based deconvolution; template time constants: 1 ms rise, 10 ms decay), and all events smaller than 4×SD of baseline noise from the highest noise recording (3.93 pA; 15.7 pA detection limit) were excluded to prevent false negative inclusion and ensure comparability across cells (following ^69^). Histogram bins start from the applied detection limit, and include an equal number of events from each cell. No correction was made for liquid junction potentials throughout.

Optogenetic experiments were performed using horizontal slices of St18-Cre:Ai9 mice injected in dorsal CA3 with a ChIEF-expressing AAV (see above). Optogenetic responses were elicited by 1 ms blue-light stimulation in 5 pulse 20 Hz trains (10 s sweeps) using a TTL-triggered CoolLED pE-300 system. Light stimulation intensity was adjusted to elicit 2–5 mV initial responses in recorded cells. All included cells were confirmed not to express channelrhodopsin by light stimulation using a 0.5 s pulse, which elicited square current responses in expressing cells. GABA_A_ and GABA_B_ receptor currents were blocked by application of 10 μM GABAzine (SR-95531; Biotrend AG) or 1 μM CGP 55845 (in DMSO; hellobio, HB0960) respectively. Excitatory synaptic transmission and feedforward nature of inhibitory responses was confirmed by excitatory receptor inhibition using 10 μM CNQX (6-cyano-7-nitroquinoxaline-2,3-dione). Response latencies were measured in voltage clamp as the time between light onset and response onset. Optogenetic-evoked potentials were recorded in current clamp configuration without current injection, and cells whose membrane potential deteriorated beyond −50 mV were excluded. Recordings which did not include both *St18+* and *St18−* cells were also excluded. The peak of PSPs was measured from the average of at least 10 sweeps per condition, and drugs were applied with the same order and timing for all experiments. Due to the different receptor onset kinetics, GABAzine-sensitive responses were calculated from the difference in PSP_3_ peak amplitude in the presence and absence of drug, while CGP 55845-sensitive responses were calculated from the most hyperpolarised potential reached within 500 ms after train stimulation. As multicellular patch-clamp data were acquired, reported cell pairs include all combinations of *St18+* and *St18−* neuron pairs from each recording.

### Immunohistochemistry and morphological analysis of patch-clamp recorded slices

After recording, slices were fixed and stained for intracellular biocytin to assess neuronal morphology. Slices were fixed with 4% PFA:PB overnight at 4°C before washing in 0.1 M PB to stop the fixation reaction. Slices were blocked by incubation with a solution containing 10% NGS in 0.1 M PB for 1–3 h at RT. Primary antibodies were added in a blocking/permeabilizing solution containing 5% NGS and 0.4% Triton X-100 overnight at RT. Slices were washed (3 × 30 min in PB) before addition of secondary antibodies and Alexa Fluor (AF) 647-conjugated streptavidin (1:333 diluted from 2 mg mL^−1^ stock solution, Invitrogen S32357) in blocking/permeabilizing solution, incubated overnight at RT with gentle shaking. Slices were washed again (3 × 10 min in PB) before clearing by incubation for 10 min at RT in CUBIC solution ^95^ (50% sucrose, 25% urea, 10% 2,2′,2″,-nitrilotriethanol, 0.1% and Triton X-100 in MilliQ water; all Sigma-Aldrich: 16104, U5128-500G, 90279-100mL), before mounting on glass slides (Assistant, Karl Hecht ref. 42406020) beneath a 1.5 thickness glass coverslip (VWR 631-0147) in CUBIC solution surrounded by a ring of Mowiol (Mowiol 4–88, Carl Roth, 713.2; Glycerol, Sigma-Aldrich, G-9012) to prevent exposure of CUBIC solution to air.

Both mouse and human recorded slices were stained with anti-NeuN (diluted 1:333 from stock; rabbit anti-NeuN, Thermo Fisher Scientific PA5-37407, RRID:AB_2554049), visualized with AF488-conjugated anti-rabbit secondary antibody (1:300 from stock; goat anti rabbit AF488, Thermo Fisher Scientific A11034, RRID:AB_2576217). Optogenetic stimulation slices were instead stained with anti-GFP (chicken polyclonal, Abcam ab13970, RRID:AB_300798) to amplify fluorescence of ChIEF-EYFP infected axons, and were visualised with AF488-conjugated anti-chicken secondary antibody (1:300 from stock; Thermo Fisher Scientific, A-11039; RRID:AB_2534096).

Slides were imaged on the ANDOR Dragonfly microscope. Overview images of all slices were taken using a 10x air objective (Nikon MRD00105, CFI P-Apo 10x, NA 0.45) and coarse (10 μm) z-steps, while cellular morphology was captured using a 20x water-immersion objective (Nikon MRD77200, CFI P-Apochromat 20x, NA 0.95), imaging all observable fluorescently labelled structures. Higher resolution images for analysis of thorny excrescences were taken using a 40x water-immersion objective (Nikon MRD77410, Apochromat LWD λS 40x, NA 1.15). Tiles were stitched using Imaris Stitcher software (Oxford Instruments). Pinhole discs with 25 or 40 μm hole diameters were used for 10x and 20x, or 40x objective images, respectively.

Stitched image stacks of recorded cells were converted to TIFF format using FIJI ^89^ for reconstruction using Neutube. Semi-automated reconstruction with manual correction of all dendritic processes was performed, producing SWC files for cells of interest. Dendritic identity was manually assigned by visual inspection. SWC files were analysed using custom MATLAB scripts. Reconstructions were corrected for compression artifacts due to mounting by scaling to slice thickness of 350 μm. Thorny excrescences were identified by visual inspection, and dendritic lengths containing thorns were manually segmented with Neutube, blind to cell type identity. Taproot dendrites were identified visually, blind to cell type identity. Shank length was measured manually in FIJI as the dendritic length from soma to the first apical branch point.

### Processing of publicly available single-cell/single-nucleus RNA-seq datasets

Single-cell RNA-seq datasets for mouse brain from Yao et al. ^41^ were used, and single-nucleus RNA-seq datasets for pig brain from Franjic et al. ^64^, and for human brain from Thompson et al. ^65^ were used. Analyses were performed in two Python environments. Doublet detection was performed in Python 3.11.8 (scanpy v1.11.0, numpy v1.26.4, pandas v2.2.2, matplotlib v3.10.1, scrublet v0.2.3). All other analyses — including data loading, quality control, dimensionality reduction, clustering, differential gene expression, and cluster stability assessment — were performed in Python 3.11.2 (scanpy v1.11.1, numpy v2.2.5, pandas v2.2.3, matplotlib v3.10.3, anndata v0.11.4, harmonypy v0.0.10, scipy v1.15.3, scikit-learn v1.5.2, seaborn v0.13.2).

Mouse hippocampal datasets (WMB-10Xv2-HPF/raw and WMB-10Xv3-HPF/raw) were downloaded in H5AD format with accompanying metadata using the abc_atlas_access Python package (Allen Brain Cell Atlas; https://github.com/AllenInstitute/abc_atlas_access), and concatenated. Pig hippocampal data and metadata were downloaded from NCBI GEO (GSE186538) and converted into H5AD format. Human hippocampal data and adjacent metadata (record EH9606; https://bioconductor.org/packages/humanHippocampus2024, humanHippocampus2024 v1.2.0) were retrieved as a SingleCellExperiment object (v1.32.0) using ExperimentHub (v3.0.0), and converted to H5AD format using anndataR (v1.0.2), in R (v4.5.3). For mouse, Ensembl gene IDs were mapped to gene symbols using the GRCm38 Ensembl release 102 GTF annotation. Quality control metrics including the percentage of mitochondrial and ribosomal gene counts were calculated per cell from raw counts for mouse and human; for pig, these metrics were assessed using pre-existing metadata from the original dataset.

All datasets were processed using a shared pipeline. Genes expressed in fewer than 10 cells were excluded, raw counts were normalized to 10,000 counts per cell and log1p-transformed. Highly variable genes (HVGs) were identified using a dataset-specific batch key (described below per species), expression values were scaled to unit variance (max=10), and PCA was performed on HVGs using the ARPACK solver. Batch effects were corrected in PCA space using Harmony (max_iter=20) ^96^; for initial exploratory clustering the *harmonypy* default diversity penalty (theta=0) was used, while all final processing steps following quality control cluster removal used theta=2. A nearest-neighbor graph (k=15) was constructed on the Harmony-corrected embedding and UMAP ^97^ was computed for visualization. Leiden clustering ^98^ was performed across a range of resolutions to identify low-quality, non-neuronal, or doublet-like clusters. Cluster identity was assessed using canonical cell type markers, QC metrics (*n_genes_by_counts*, *total_counts*, percentage of mitochondrial and ribosomal counts), UMAP topology, and Scrublet doublet prediction scores ^99^ where applicable. Following removal of low-quality clusters, HVG identification, scaling, PCA, and Harmony correction (theta=2) were repeated on the cleaned dataset. This iterative cleaning and reprocessing approach was applied independently at each level of subsetting.

For mouse, cells were subset to DG GCs (subclass ‘037 DG Glut’), immature GCs (supertype ‘0141 DG-PIR Ex IMN_2’), CA1 PNs (subclass ‘016 CA1-ProS Glut’), CA2 PNs (subclass ‘025 CA2-FC-IG Glut’), CA3 PNs and mossy cells (subclass ‘017 CA3 Glut’), retaining 119,675 cells and 25,820 genes. 2,361 HVGs were identified using the library version (’library_method’) as the batch key, and UMAP was computed using 45 Harmony-corrected PCs. No low-quality clusters required removal at the hippocampal formation level. The CA3 and CA1 subsets were independently derived from the full concatenated dataset and the same preprocessing pipeline described above was applied to both. For CA3, cells were subset using supertypes ‘0075 CA3 Glut_1’, ‘0076 CA3 Glut_2’, ‘0077 CA3 Glut_3’, and ‘0078 CA3 Glut_4’, retaining 4,536 cells, 20,817 genes, and 2,215 HVGs, and clustered with 20 Harmony-corrected PCs. For CA1, cells were subset using subclass ‘016 CA1-ProS Glut’, retaining 31,336 cells, 24,240 genes, and 2,126 HVGs, and clustered with 30 Harmony-corrected PCs.

For pig, nuclei were subset to annotated clusters ‘GC’, ‘CA1 Sub’, ‘CA2-3’, and ‘MC’, retaining 7,672 nuclei and 23,058 genes. 2,000 HVGs were identified using sample name (’samplename’) as the batch key, and Leiden clustering was performed using 20 Harmony-corrected PCs. At Leiden resolution 0.58, one cluster (125 nuclei) was removed based on convergent evidence: isolated UMAP topology, low *n_genes_by_counts* and *total_counts*, absence of *Slc17a7* expression, and a 36% predicted doublet rate obtained with Scrublet. Three additional clusters showed oligodendrocyte marker enrichment (*Mobp, Mbp, Plp1, and Opalin)* but were conservatively retained due to topological embedding within larger neuronal populations. Following cluster removal and reprocessing, a final hippocampal dataset of 7,547 nuclei was retained with 20 Harmony-corrected PCs for UMAP. The pig CA2-3 subset was derived from the ‘CA2-3’ cluster (1,756 nuclei) from the final hippocampal dataset, and after Leiden clustering using 10 Harmony-corrected PCs, a cluster of 99 nuclei enriched for oligodendrocyte markers was removed at resolution 0.16, retaining a final dataset of 1,657 nuclei with 10 PCs for UMAP. The pig CA3 subset was defined as *Rgs14*-negative nuclei at Leiden resolution 0.02 from the final CA2-3 dataset (i.e. removing CA2 cells), retaining 1,531 nuclei with 10 PCs for UMAP.

For human, nuclei were subset to ‘GC’, ‘CA2-4’, and ‘CA1/ProS’ based on the ‘fine.cell.class’ label, retaining 20,320 nuclei and 31,451 genes. 2,000 HVGs were identified using donor ID (’brnum’) as the batch key, and Leiden clustering was performed using 45 Harmony-corrected PCs. At Leiden resolution 0.6, four clusters totaling 715 nuclei were removed: one showed enrichment for oligodendrocyte and astrocyte markers (*MBP*, *MOBP*, *PLP1*, *GFAP*), one showed elevated *GAD2* and *RELN* expression associated with Cajal-Retzius cells, one showed elevated expression of *PLP1* and *GFAP* and non-neuronal marker *MECOM*, and one presented a 42.9% predicted doublet rate by Scrublet. Following cluster removal and reprocessing with 40 Harmony-corrected PCs, at Leiden resolution 0.7 a further cluster of 328 nuclei with low *n_genes_by_counts* and *total_counts* was removed, obtaining a final human hippocampal dataset of 19,277 nuclei, with 45 Harmony-corrected PCs used for UMAP. The human CA2-3 dataset was derived by subsetting the final human hippocampal dataset using ‘CA2’, ‘CA3.1’ and ‘CA3.2’ labels from ‘superfine.cell.class’; donor Br2743 (1 nucleus) was excluded, yielding 4,147 nuclei. After Leiden clustering using 15 Harmony-corrected PCs, a topologically isolated cluster of 26 nuclei was identified; differential expression analysis using the Wilcoxon rank-sum test (LINC-genes excluded) revealed enrichment for ribosomal genes, and the cluster was removed, retaining a final CA2-3 dataset of 4,121 nuclei with 10 PCs for UMAP. The human CA3 dataset was derived by subsetting ‘CA3.1’ and ‘CA3.2’ from the final CA2-3 dataset, retaining 3,745 nuclei with 10 PCs for UMAP.

### Differentially expressed genes and cluster stability analysis

Differentially expressed genes (DEGs) were identified across all species using a Wilcoxon rank-sum test on log-normalized expression values, comparing each cluster against all others. Marker genes were retained based on the following criteria: Benjamini–Hochberg-adjusted p-value < 0.05, expression in > 80% of cells within the cluster, and a specificity score > 0.3 (defined as the difference in non-zero expression fraction between the cluster and the reference population). The log2 fold-change threshold was set to > 2 for mouse and human, and relaxed to > 1.5 for pig to ensure a sufficient number of marker genes could be identified per cluster. Prior to DEG analysis, genes lacking conserved symbols were excluded to facilitate cross-species comparison (pig genes prefixed “LOC”, and human long intergenic non-coding RNAs prefixed “LINC”).

To visualize cluster stability across Leiden resolutions, a Sankey diagram was constructed in SankeyMATIC (https://sankeymatic.com/) using the transitions of cells between clustering resolutions, selecting resolutions that first introduced each new cluster. Clustering stability was assessed by computing the Adjusted Rand Index (ARI; a measure of similarity between two clusterings corrected for chance, where 1 indicates identical partitions and 0 indicates random agreement; *adjusted_rand_score* from scikit-learn) between consecutive Leiden clustering solutions, and silhouette scores (a measure of how well each cell fits its assigned cluster relative to neighbouring clusters, ranging from −1 to 1 with higher values indicating better-defined clusters; *silhouette_score* from scikit-learn) on the Harmony-corrected PCA embedding across tested resolutions.

### Spatial transcriptomic analysis

Mouse spatial transcriptomic data were presented using the Allen ABC Atlas. Deep and superficial PN locations (**Fig. 1b**) were acquired by cross-referencing Allen ABC cell ‘cluster’ identities to Leiden clusters 1 and 2 from our analysis (see **Supplementary Fig. 1d**). Allen clusters 0296-0301 and 0304-0306 were assigned as deep *St18+* PNs (our cluster 2), while Allen clusters 0302-0303 and 0307-0315 were assigned as superficial *St18−* PNs (our cluster 1). *St18* spatial distribution presents expression levels across all CA3 PNs (supertypes: 0075 CA3 Glut_1, 0076 CA3 Glut_2, 0077 CA3 Glut_3, 0078 CA3 Glut_4), excluding mossy cells (supertype 0079 CA3 Glut_5).

Human spatial transcriptomic data were acquired and replotted from Thompson et al. ^65^. Sample Br8667 was analysed, with cell type localisation presented by plotting cell-specific NMF signature values normalised within each condition (DG GCs, sum of nmf5, 10, 14, and 26; mossy cells, nmf52; sup CA3, nmf11; deep CA3, nmf63; CA2, nmf61; CA1, nmf15). A 2-mm-thick line profile was used to visualise deep-superficial locations of CA3 subtypes, measuring peak normalised NMF scores using a 50 µm bin-width.

### Quantification and statistical analysis

All data are presented and reported as median (with interquartile range, IQR) unless otherwise stated. Box plots depict median (line), IQR (box) and minimum to maximum values (whiskers) with individual datapoints overlain. Where used, symbols with error bars represent mean ± SEM. Unpaired datasets were compared using non-parametric statistical tests for either two-sample (Mann-Whitney test) or multi-sample (Kruskal-Wallis test with Dunn’s multiple comparisons test) data unless stated otherwise in the figure legend. Paired datasets were compared using a Wilcoxon signed-rank test. All other tests are reported in figure legends where used. All tests were two-sided and exact p-values are reported throughout (significance reported at *p* < 0.05). ‘n’ refers to cells or pairs and ‘N’ to animals or patients as specified throughout. Statistical analysis was performed using Graphpad Prism (v11).

## DATA AND CODE AVAILABILITY

All RNA sequencing datasets analysed in this study are publicly available. Mouse hippocampal data were obtained from the Allen Brain Cell Atlas ^41,43^ (https://github.com/AllenInstitute/abc_atlas_access). Pig hippocampal data ^64^ are available from NCBI GEO (GSE186538). Human hippocampal data ^65^ are available through an ExperimentHub Data package (https://bioconductor.org/packages/humanHippocampus2024). Mouse spatial transcriptomic data were accessed using the Allen Brain Cell Atlas online platform (https://brain-map.org/bkp/explore/abc-atlas), while human spatial transcriptomic data were acquired via the humanHippocampus2024 Bioconductor package (https://bioconductor.org/packages/release/data/experiment/html/humanHippocampus2024.html) using the samui browser (https://samuibrowser.com/from?url=data.libd.org/samuibrowser/&s=Br8667). All other data supporting the findings of this study are available from the corresponding authors upon request. Custom code used for patch-clamp data analysis is publicly available at https://github.com/jakefwatson/patchanalysis. Scripts for bioinformatics analysis are available on request.

## ACKNOWLEDGEMENTS

We are very grateful to Liset M. de la Prida for critical feedback on the manuscript. We thank Florian Marr, Christina Altmutter and Marion Ederer for excellent technical support, Eleftheria Kralli-Beller for manuscript editing, Alois Schlögl for computational assistance, and Andrea Navas-Olive for coding support and comments on the manuscript. This research was supported by the Scientific Services Units (SSUs) of ISTA. We thank Victoria Wimmer, Mary Muhia, and the Preclinical Facility for excellent animal management, Daniel Spies, Andrea Garofoli, and Scientific Computing for support with bioinformatic analysis, Todor Asenov and the Miba Machine Shop, the Imaging and Optics Facility, the Lab Support Facility, the Virus Service and Fangfang Sun for outstanding support with viral vector production, and Alazne Dominguez Monedero for support with single-cell analysis. We are deeply grateful to the patient donors for their consent to contribute to this project, and acknowledge the excellent support of the Medical University of Vienna Department of Neurosurgery staff; Romana Hoeftberger and the Division of Neuropathology and Neurochemistry; Gregor Kasprian and the Division of Neuroradiology and Musculoskeletal Radiology; Matthias Tomschik for assistance with the ethical application; and Christoph Baumgartner, Martha Feucht, and Ekaterina Pataraia for their clinical care of the patients included in this study. We thank Rosa Cossart for *Ngn2*-CreER mice and several insightful discussions on developmental neuronal labelling. The project received funding from the European Research Council (ERC) under the European Union’s Horizon 2020 research and innovation programme (advanced grant No 692692 GIANTSYN and 101199096 CA3-SYNGRAM to P.J., and Marie Skłodowska-Curie Actions Individual Fellowship no. 101026635 to J.F.W.) and the Austrian Science Fund (FWF; grant PAT 4178023 and 10.55776/CoE16 to P.J.).

## AUTHOR CONTRIBUTIONS

J.F.W. and P.J. conceived the study. J.F.W. and R.J.M.M. designed and performed all experiments. R.J.M.M. performed bioinformatic analysis. Y.B.S. produced viruses and advised on viral tracing experiments. S.A.J. generated the transgenic *St18*-Cre mouse line. K.R. performed neurosurgery. J.F.W. and P.J. supervised the study and acquired funding. J.F.W. wrote the paper and all authors revised the manuscript.

