## Supplementary Information for "Hippocampal CA3 forms a two-layer network of molecularly distinct cell types in mice and humans"

### Supplementary Information for Morse-Mora et al.

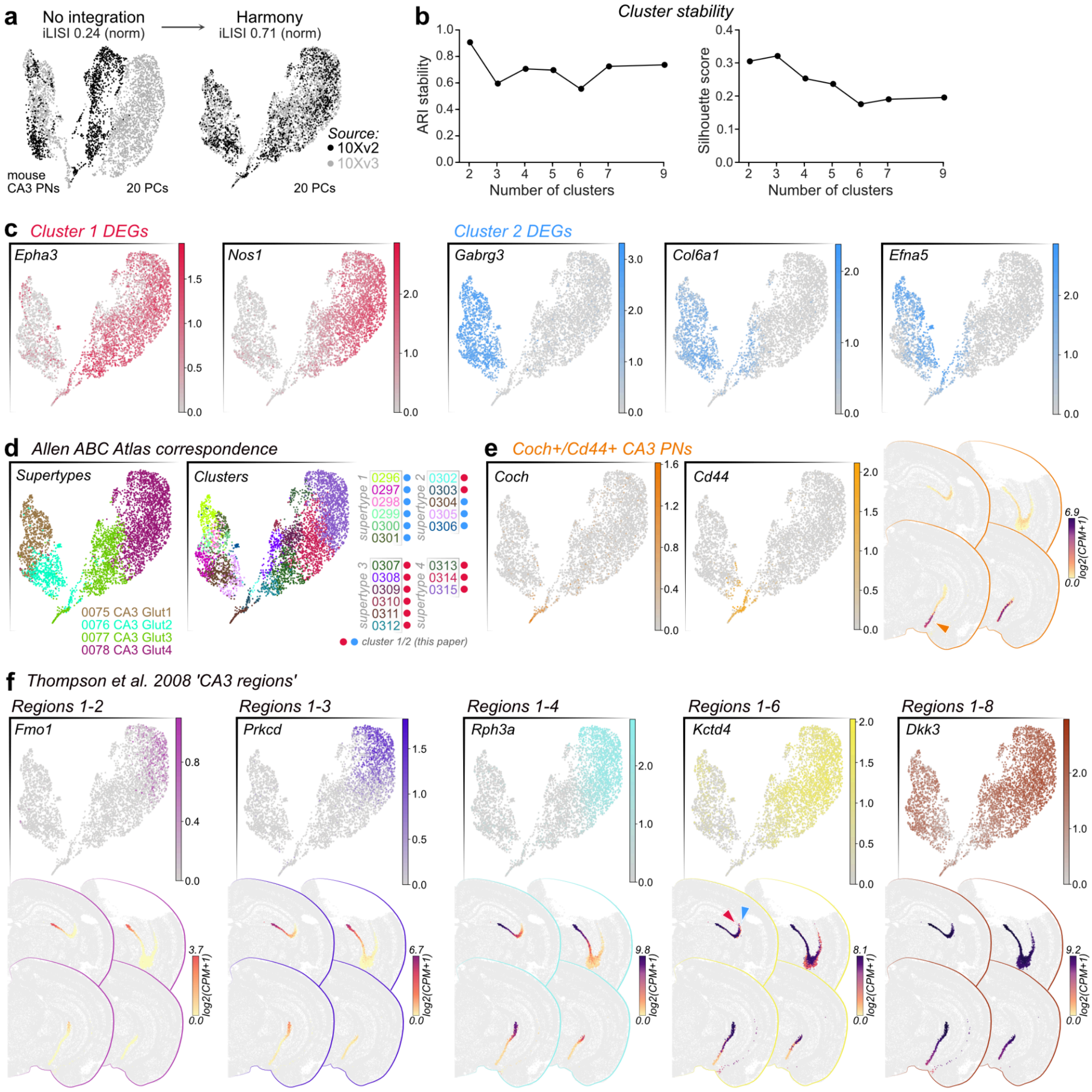


**Supplementary Figure 1: Transcriptomic analysis of CA3 PN heterogeneity.**
**a**, UMAP representation of CA3 PN scRNA-seq data from Yao et al. 2023 [41] 10Xv2 and 10Xv3 datasets pre and post harmonisation. **b**, Measures of cluster stability for CA3 PN Leiden clusters (see **Fig. 1**) show high ARI stability (Adjusted Rand Index) and silhouette score for low cluster numbers. **c**, Example expression distributions of differentially expressed genes for cluster 1 (superficial; red) and 2 (deep; blue) (units: ln(CP10k+1) expression). **d**, Overlaying Allen ABC Atlas supertypes and clusters on our CA3 PN heterogeneity distribution shows that supertypes 0077 and 0078 map largely to the superficial PN subtype, while 0075 and 0076 map to the deep PN subtype. ABC clusters 0302 and 0303 do not fit this principle, both forming part of the superficial PN subtype in our dataset. Circles represent superficial (cluster 1) and deep (cluster 2) identity from this study, and cluster assignment used for spatial transcriptomics in **Fig. 1b**. **e**, *Coch* and *Cd44* (expression units: ln(CP10k+1)) mark a small subpopulation of CA3 PNs located in the very ventral hippocampus (arrowhead), seen on ABC atlas spatial transcriptomic data for *Coch*. **f**, Marker genes (UMAP expression units: ln(CP10k+1)) for CA3 'regions' reported in Thompson et al. 2008 [24] sweep across the superficial PN cluster, with no labelling of the deep PN cluster except when CA3 is completely labelled (*Dkk3*). *Fmo1*, *Prkcd*, and *Rpha3* show strong proximal-distal expression gradients in spatial transcriptomic data, while clear preferential labelling of superficial but not deep PNs is evident with *Kctd4* (arrowheads indicate superficial, red; and deep, blue).

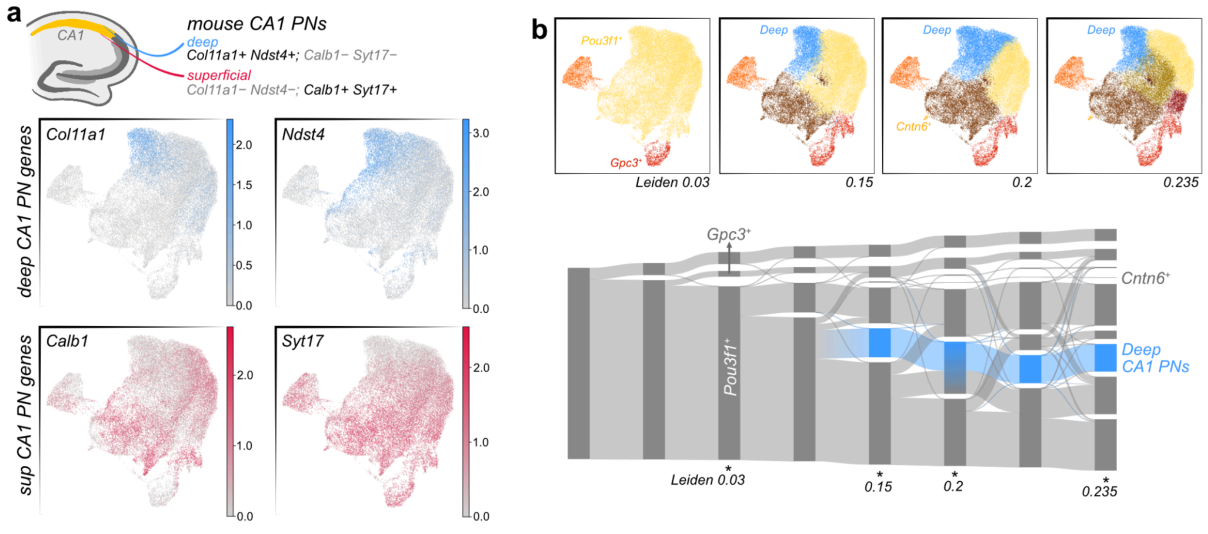


**Supplementary Figure 2: Transcriptomic analysis of CA1 PN heterogeneity.**

**a**, Classical markers of deep CA1 PNs allow identification of their location on UMAP plots. Mouse CA1 PNs from Yao et al. 2023 are presented (expression units: ln(CP10k+1)). **b**, Leiden clustering of CA1 does not segregate deep and superficial PNs at low resolution, instead, deep PNs emerge as an unstable cluster at higher resolution, consistent with deep-superficial heterogeneity forming a gradient rather than distinct cell types in CA1. Markers of select clusters are highlighted. Neurons consistently comprising the deep CA1 PN cluster are highlighted on the Sankey plot.

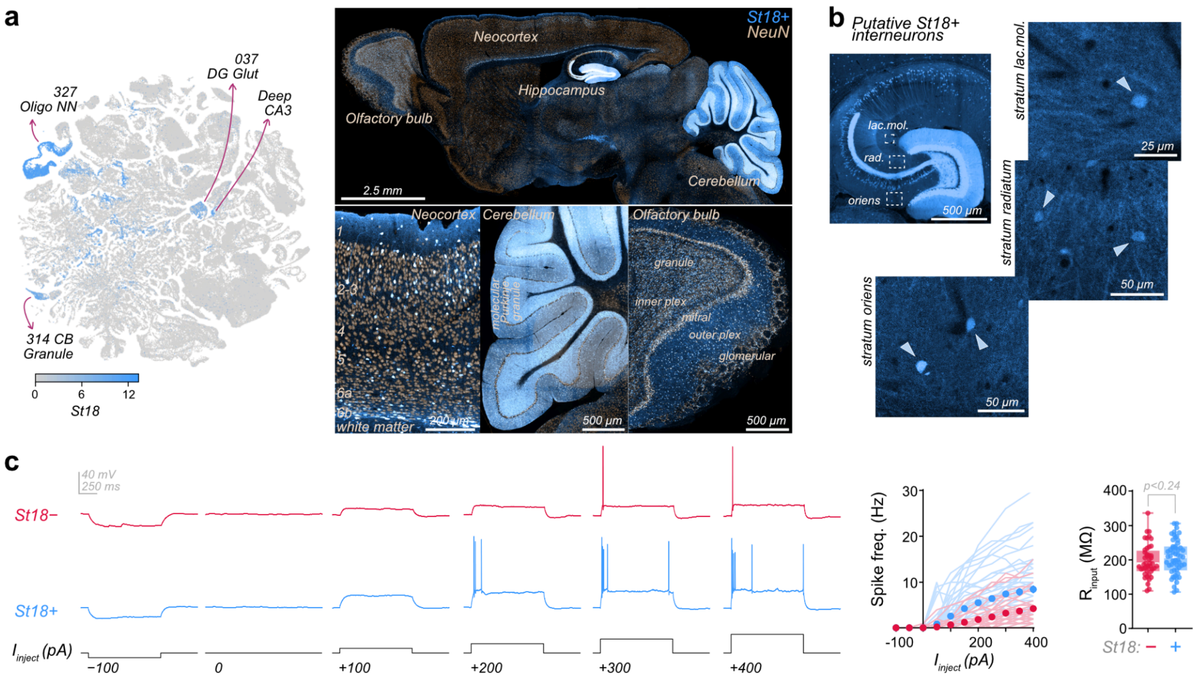


**Supplementary Figure 3: Distribution of labelled cells in *St18*-Cre mice.**

**a**, *St18* expression in Allen Brain Cell Atlas RNA-seq data shows labelling of cell populations beyond the hippocampus, with equivalent labelling seen in *St18*-Cre mice. **b**, Labelling of putative interneurons by *St18*-Cre:Ai9 in hippocampal CA3 *stratum oriens*, *radiatum*, and *lacunosum moleculare*. **c**, Example traces of *St18*+ and *St18*− CA3 PN firing responses upon increasing current injection, with quantified spike frequency across 1-s pulses (symbols present mean ± SEM), and quantified input resistance (*St18*−, 188 MΩ (169, 227), *n* = 44; *St18*+, 209 MΩ (170, 239), *n* = 54; Mann-Whitney test, *p* = 0.24).

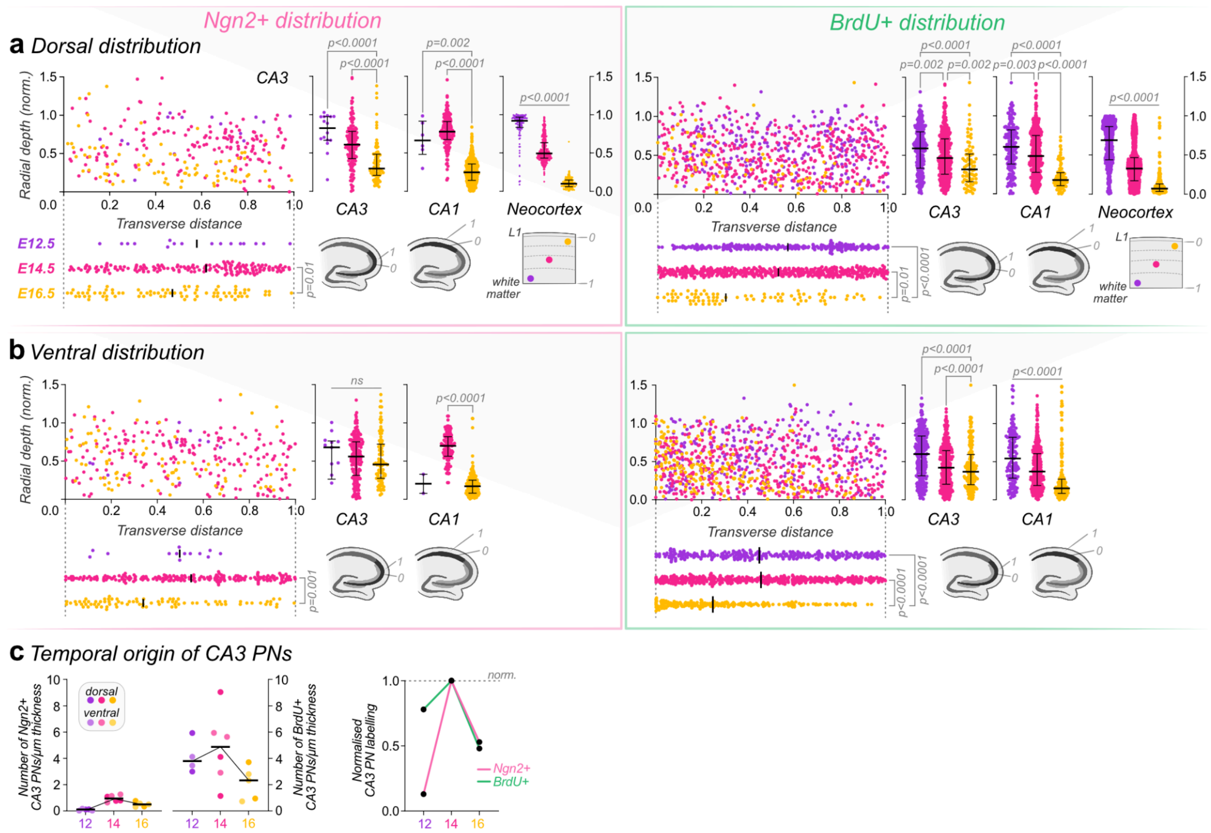


**Supplementary Figure 4: *Ngn2* and BrdU birthdating recapitulate the same distribution and temporal patterns.**

**a**, Dorsal and **b,** ventral distribution of *Ngn2*-CreER labelled and BrdU+ neurons in CA3 and CA1 follow the same deep-to-superficial radial distribution, while neurons in the neocortex follow a white matter-to-layer I gradient from E12.5 to E16.5. Both labelling strategies also show E16.5-born CA3 PNs are positioned more proximal than earlier-born neurons (line represents median, and error bars denote IQR. For summary data see **Supplementary Table 1**). All statistical comparisons were performed with a Kruskal-Wallis test. **c**, The density of *Ngn2*-labelled CA3 PNs (per µm of tissue thickness) is ~4 fold lower than that of BrdU+ CA3 PNs (*Ngn2*: E12.5, 0.12 (0.03, 0.16), *N* = 4, 4 TMs; E14.5, 0.94 (0.74, 1.2), *N* = 3, 3 TMs; E16.5, 0.5 (0.33, 0.67), *N* = 3, 3 TMs. BrdU: E12.5, 3.8 (3.1, 5.5), *N* = 2, 2 TMs; E14.5, 4.9 (2.5, 6.7), *N* = 3, 3 TMs; E16.5, 2.3 (0.84, 3.3), *N* = 3, 3 TMs; line represents median); nonetheless, both birthdating strategies recapitulate the same overall pattern, with CA3 PN generation peaking at E14.5 (E14.5 normalized to 1.0; *Ngn2* E12.5: 0.13; E16.5: 0.53; BrdU E12.5: 0.78; E16.5: 0.48).

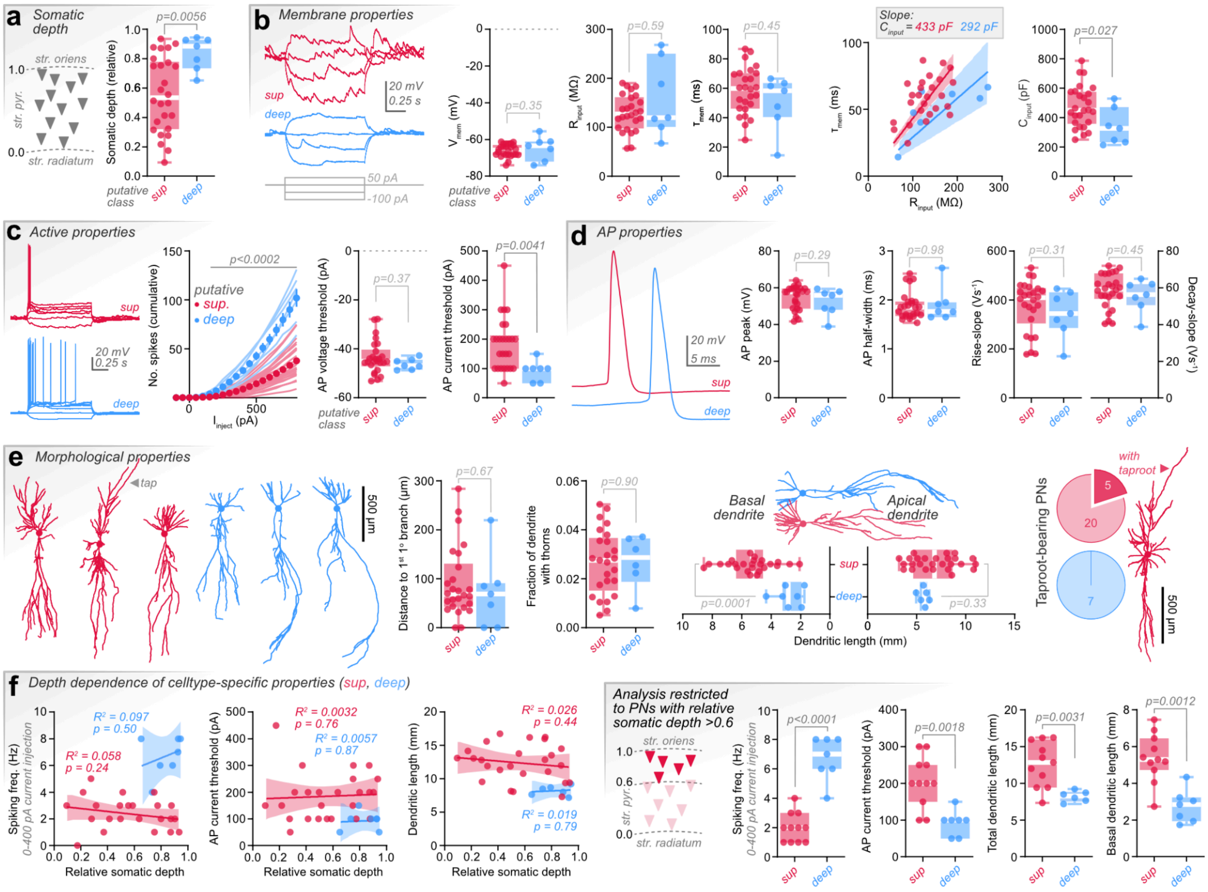


**Supplementary Figure 5: Distinct morpho-functional properties of putative deep and superficial CA3 PNs in humans.**

**a**, Relative somatic location of putatively classified CA3 PNs (median (IQR): sup, 0.53 (0.32, 0.78), *n* = 26; deep, 0.89 (0.74, 0.93), *n* = 7; Mann-Whitney test, *p* = 0.0056). **b**, Passive membrane properties are similar between PN types, differing only in cell capacitance (*n* = 26 sup and *n* = 7 deep PNs throughout. $V_{mem}$: sup, −65.8 mV (−68.3, −62.9); deep, −63.4 mV (−72.0, −61.1); *p* = 0.35. $R_{input}$: sup, 127.5 MΩ (98.2, 161.5); deep, 120.0 MΩ (101.1, 250.8); *p* = 0.59. $\tau_{mem}$: sup, 59.9 ms (46.2, 70.2); deep, 58.4 ms (40.4, 63.4); *p* = 0.45. $C_{input}$: sup, 450.3 pF (357.7, 575.8); deep, 326.6 pF (232.8, 470.2); *p* = 0.027). **c**, Putative deep PNs show higher firing (AP: action potential) and lower rheobase than sup PNs (*n* = 26 sup and *n* = 7 deep PNs throughout. Cumulative spikes upon increasing current injection, *p* < 0.0002 from 150 pA injection, multiple Mann-Whitney tests with Bonferroni-Dunn correction. AP voltage threshold: sup, −45.1 mV (−47.1, −40.3); deep, −46.9 mV (−48.5, −44.4); *p* = 0.37. AP current threshold: sup, 200 pA (100, 212.5); deep, 100 pA (50, 100); *p* = 0.0041). **d**, No difference in AP properties was observed between PN types (*n* = 26 sup and *n* = 7 deep PNs throughout. AP peak from 0 mV: sup, 57.7 mV (48.7, 59.6); deep, 55.8 mV (48.0, 57.9); *p* = 0.29. AP half-width: sup, 1.85 ms (1.67, 1.97); deep, 1.77 ms (1.65, 1.95); *p* = 0.98. AP rise slope: sup, 409.4 $V \cdot s^{-1}$ (304.5, 448.6); deep, 344.7 $V \cdot s^{-1}$ (286.4, 431.9); *p* = 0.31. AP decay slope: sup, 59.3 $V \cdot s^{-1}$ (54, 67.8); deep, 56.3 $V \cdot s^{-1}$ (50.7, 62.1); *p* = 0.45). **e**, Human deep PNs lack some subtype-characteristic properties seen in mice (Distance from soma to first primary branch point: sup, 68.7 μm (42.9, 131.1), *n* = 26; deep, 69.1 μm (0.0, 90.8), *n* = 7; *p* = 0.67. Fraction of total dendritic length bearing thorns: sup, 0.028 (0.015, 0.037), *n* = 25; deep, 0.029 (0.019, 0.037), *n* = 6; *p* = 0.90) and have simpler branching morphology than superficial PNs (basal dendritic length: sup, 5.3 mm (4.6, 6.4), *n* = 25; deep, 2.8 mm (1.9, 3.2), *n* = 7; *p* = 0.0001. Apical dendritic length: sup, 6.8 mm (4.7, 8.7), *n* = 26; deep, 5.8 mm (5.3, 6.1), *n* = 6; *p* = 0.33). Taproot-bearing CA3 PNs were exclusively of superficial cell type in this dataset (5/20 sup cells, 0/7 deep cells). **f**, No putative 'cell type-specific properties' showed relation to somatic depth within classes. These observations are consistent with distinct subtypes, rather than a model where CA3 PNs form a heterogeneous gradient (linear regressions with $R^2$ displayed, p-values indicate whether regression slope is significantly non-zero). Similarly, comparing properties of deep PNs and sup PNs with relative somatic depth >0.6 confirms cell type-specific properties are not a result of somatic depth (spiking frequency: sup, 2 Hz (1, 3), *n* = 11; deep, 7 Hz (6, 8), *n* = 7; *p* < 0.0001. AP current threshold: sup, 200 pA (150, 250), *n* = 11; deep, 100 pA (50, 100), *n* = 7; *p* = 0.0018. Total dendritic length: sup, 12.9 mm (9.4, 16.0), *n* = 11; deep, 8.3 mm (7.5, 8.8), *n* = 6; *p* = 0.0031. Basal dendritic length: sup, 5.3 mm (4.7, 6.5), *n* = 11; deep, 2.8 mm (1.9, 3.2), *n* = 7; *p* = 0.0012).

**Supplementary Table 1: Statistics of Ngn2-CreER and BrdU cell labelling data**
Summary of birthdating data where *n* denotes cells, *N* denotes animals, TM denotes timed-matings.

| ***Ngn2-CreER labelling*** | | | | | | |
|---|---|---|---|---|---|---|
| | **E12.5** | | **E14.5** | | **E16.5** | |
| | **Median (IQR)** | ***n* (*N*, TMs)** | **Median (IQR)** | ***n* (*N*, TMs)** | **Median (IQR)** | ***n* (*N*, TMs)** |
| *Transverse distance (normalised)* | | | | | | |
| **CA3 dorsal** | 0.58 (0.34, 0.78) | 16 (2, 2) | 0.62 (0.34, 0.79) | 168 (3, 3) | 0.47 (0.19, 0.69) | 82 (2, 2) |
| **CA3 ventral** | 0.5 (0.32, 0.6) | 13 (3, 3) | 0.55 (0.28, 0.78) | 202 (3, 3) | 0.34 (0.17, 0.62) | 84 (3, 3) |
| *Radial depth (normalised)* | | | | | | |
| **CA3 dorsal** | 0.82 (0.67, 0.97) | 16 (2, 2) | 0.61 (0.43, 0.78) | 168 (3, 3) | 0.3 (0.21, 0.48) | 82 (2, 2) |
| **CA3 ventral** | 0.68 (0.26, 0.76) | 13 (3, 3) | 0.56 (0.31, 0.75) | 202 (3, 3) | 0.46 (0.28, 0.72) | 84 (3, 3) |
| **CA1 dorsal** | 0.66 (0.48, 0.91) | 6 (2, 2) | 0.78 (0.66, 0.91) | 144 (3, 3) | 0.25 (0.14, 0.36) | 477 (2, 2) |
| **CA1 ventral** | 0.2 (0.08, 0.33) | 2 (3, 3) | 0.7 (0.56, 0.82) | 119 (3, 3) | 0.17 (0.08, 0.25) | 239 (3, 3) |
| **Neocortex** | 0.92 (0.83, 0.97) | 118 (2, 2) | 0.49 (0.43, 0.64) | 585 (3, 3) | 0.1 (0.06, 0.15) | 178 (2, 2) |
| ***BrdU labelling*** | | | | | | |
| | **E12.5** | | **E14.5** | | **E16.5** | |
| | **Median (IQR)** | ***n* (*N*, TMs)** | **Median (IQR)** | ***n* (*N*, TMs)** | **Median (IQR)** | ***n* (*N*, TMs)** |
| *Transverse distance (normalised)* | | | | | | |
| **CA3 dorsal** | 0.57 (0.33, 0.78) | 289 (2, 2) | 0.52 (0.26, 0.77) | 451 (3, 3) | 0.3 (0.17, 0.62) | 83 (2, 2) |
| **CA3 ventral** | 0.45 (0.24, 0.72) | 295 (2, 2) | 0.46 (0.2, 0.69) | 486 (3, 3) | 0.25 (0.09, 0.47) | 219 (3, 3) |
| *Radial depth (normalised)* | | | | | | |
| **CA3 dorsal** | 0.59 (0.33, 0.8) | 289 (2, 2) | 0.47 (0.26, 0.71) | 451 (3, 3) | 0.32 (0.16, 0.52) | 83 (2, 2) |
| **CA3 ventral** | 0.6 (0.31, 0.83) | 295 (2, 2) | 0.42 (0.2, 0.64) | 486 (3, 3) | 0.37 (0.2, 0.59) | 219 (3, 3) |
| **CA1 dorsal** | 0.6 (0.38, 0.82) | 182 (2, 2) | 0.48 (0.27, 0.75) | 545 (3, 3) | 0.18 (0.11, 0.27) | 118 (2, 2) |
| **CA1 ventral** | 0.54 (0.28, 0.82) | 146 (2, 2) | 0.37 (0.19, 0.61) | 415 (3, 3) | 0.15 (0.08, 0.27) | 220 (3, 3) |
| **Neocortex** | 0.69 (0.44, 0.86) | 1090 (2, 2) | 0.33 (0.17, 0.47) | 1903 (3, 3) | 0.07 (0.03, 0.13) | 274 (2, 2) |
